# Discovery of A New Class of Integrin Antibodies for Fibrosis

**DOI:** 10.1101/2020.07.20.207555

**Authors:** Ji Zhang, Tao Wang, Ashmita Saigal, Josephine Johnson, Jennifer Morrisson, Sahba Tabrizifard, Scott A. Hollingsworth, Michael J. Eddins, Wenxian Mao, Kim O’Neill, Margarita Garcia-Calvo, Ester Carballo-Jane, DingGang Liu, Taewon Ham, Qiong Zhou, Weifeng Dong, Hsien-Wei Yvonne Meng, Jacqueline Hicks, Tian-Quan Cai, Taro Akiyama, Shirly Pinto, Alan C. Cheng, Thomas Greshock, John C. Marquis, Zhao Ren, Saswata Talukdar, Hussam Hisham Shaheen, Masahisa Handa

**Affiliations:** Departments of Cardiometabolic Diseases, MRL, Merck & Co., Inc., 2000 Galloping Hill Road, Kenilworth, NJ 07033, USA; Departments of Discovery Biologics, MRL, Merck & Co., Inc., 2000 Galloping Hill Road, Kenilworth, NJ 07033, USA; Departments of Quantitative Biosciences, MRL, Merck & Co., Inc., 2000 Galloping Hill Road, Kenilworth, NJ 07033, USA; Departments of Computational & Structural Chemistry, MRL, Merck & Co., Inc., 2000 Galloping Hill Road, Kenilworth, NJ 07033, USA; Departments of in vitro Pharmacology, MRL, Merck & Co., Inc., 2000 Galloping Hill Road, Kenilworth, NJ 07033, USA; Departments of SALAR, MRL, Merck & Co., Inc., 2000 Galloping Hill Road, Kenilworth, NJ 07033, USA; Departments of Chemistry, MRL, Merck & Co., Inc., 2000 Galloping Hill Road, Kenilworth, NJ 07033, USA; Departments of in vivo Pharmacology, MRL, Merck & Co., Inc., 2000 Galloping Hill Road, Kenilworth, NJ 07033, USA

## Abstract

Lung fibrosis, or the scarring of the lung, is a devastating disease with huge unmet medical need. There are limited treatment options and its prognosis is worse than most types of cancer. We previously discovered that MK-0429 is an equipotent pan-inhibitor of all αv integrins that reduces proteinuria and kidney fibrosis in a preclinical model. In the present study, we further demonstrated that MK-0429 significantly inhibits fibrosis progression in a bleomycin-induced lung injury model. In search of newer integrin inhibitors for fibrosis, we characterized monoclonal antibodies discovered using Adimab’s yeast display platform. We identified several potent neutralizing integrin antibodies with unique human and mouse cross-reactivity. Among these, Ab-31 blocked the binding of multiple αv integrins to their ligands with IC50s comparable to those of MK-0429. Furthermore, both MK-0429 and Ab-31 suppressed integrin-mediated cell adhesion and latent TGFβ activation. In IPF patient lung fibroblasts, TGFβ treatment induced profound αSMA expression in phenotypic imaging assays and Ab-31 demonstrated superior in vitro activity at inhibiting αSMA expression, suggesting that the integrin antibody is able to modulate TGFβ action though mechanisms beyond the inhibition of latent TGFβ activation. Together, our results highlight the potential to develop newer integrin therapeutics for the treatment of fibrotic lung diseases.

**One Sentence Summary:** targeting integrin in lung fibrosis

## Introduction

Idiopathic pulmonary fibrosis (IPF) is a chronic, fibrosing interstitial lung disease with unknown etiology. Patients suffer from chronic coughs and deteriorating breathing difficulties. The median survival is 2.5-3.5 years from diagnosis. Despite the severe clinical impact, there are limited treatment options for lung fibrosis. In 2014, the FDA approved the use of Pirfenidone and Nintedanib in IPF patients. Both drugs slow the decline of lung function as measured by the decrease of FVC (forced vital capacity), a surrogate endpoint measurement ^1^. However, neither drug appears to stop disease progression, relieve breathing difficulty, or substantially improve patient survival. There is an unmet medical need to develop new IPF therapies that bring clinically meaningful efficacy to patients.

In recent years, the integrin family of cell adhesion molecules has emerged as key mediators of tissue fibrosis. Among the 24 known integrin heterodimers, five αv integrins (αvβ1, αvβ3, αvβ5, αvβ6, and αvβ8) transduce mechanical and biochemical signals from fibrotic extracellular matrix into the cell, activate latent TGFβ, and subsequently modulate fibroblast adhesion, migration, and growth ^2^. The αv integrins primarily interact with the RGD (Arginine-Glycine-Aspartic acid) peptide present in fibronectin and vitronectin (αvβ1, αvβ3, and αvβ5), or with the RGD motif of the TGFβ latency–associated peptide (LAP) (αvβ1, αvβ6, and αvβ8) ^2–5^. As a result, αv integrins play a key role in the regulation of TGFβ signaling ^6^. Dysregulated expression and response to TGFβ has been implicated in a wide variety of disease processes including fibrotic disease and chronic inflammation ^7^. The epithelium-specific αvβ6 integrin binds to latent TGFβ and facilitates release of the mature cytokine, a process called TGFβ activation ^3,8^. Deletion of β6 integrin *in* mice is protective against bleomycin-induced lung fibrosis ^3^, and an anti-mouse αvβ6 antibody has shown similar beneficial effects in preclinical animal studies ^9^. An αvβ6 antibody (BG00011/Biogen) and a small molecule inhibitor GSK3008348 were used in clinical trials of IPF patients (clinicaltrials.gov identifier NCT03573505, NCT03069989) ^10^. αvβ1, the less-known member of the integrin family, was recently shown to be highly expressed in activated fibroblasts and modulate lung and liver fibrosis in mice ^5^. Additionally, αvβ8 integrin, another regulator of latent TGFβ activation, modulates chemokine secretion and dendritic cell trafficking ^4,11^. β8 knockout mice and mice treated with blocking antibody are protected against airway inflammation and fibrosis ^4,12^. Although the role of a pan-αv inhibitor has not been extensively tested in the clinic for lung indications, evidence from multiple lines of work suggest that modulating αv integrin activity will lead to anti-fibrotic effects in various tissues. A report by Henderson *et al*. demonstrated that depletion of αv integrin in myofibroblast cells lead to protection against hepatic fibrosis induced by carbon tetrachloride, renal fibrosis induced by unilateral ureter obstruction, and lung fibrosis induced by bleomycin ^6^. Furthermore, a small molecule RGD mimetic CWHM12 similarly attenuates liver and lung fibrosis ^6^. The complexity of the integrins and their role in the progression of the disease suggest that a pharmacological inhibitor of multiple integrin subtypes would be required to produce meaningful effects on delaying or inhibiting the progression of fibrosis. Interestingly, recent genome-wide association analysis of 400,102 individuals identifies a strong correlation of decrease in αv integrin expression with lung function decline (FEV1/FVC), indicating a beneficial effect of αv inhibition in broader patient populations, such as chronic obstructive pulmonary disease (COPD) ^13^.

Historically, MSD has contributed significantly to the development of integrin therapeutics by bringing forth the first approved small molecule inhibitor Aggrastat (Tirofiban) for acute coronary artery syndrome ^14, 15^. We also developed a small molecule integrin inhibitor MK-0429 with good oral bioavailability in humans ^16^. MK-0429 was initially designed as an RGD mimetic against αvβ3 integrin that incorporates key pharmacophores representing the guanidine and carboxylic acid of the RGD tripeptide sequence ^17,18^. We recently found that MK-0429 is an equipotent pan-inhibitor of all αv integrins, and it reduces proteinuria and renal fibrosis in an experimental diabetic nephropathy model ^19^. In the present work, we further demonstrate that MK-0429 significantly inhibits fibrosis progression in a bleomycin-induced lung fibrosis mouse model. We also identified several potent αv integrin antibodies with unique human and mouse cross-species affinity. Among these, Ab-31 demonstrated superior in vitro activity at inhibiting TGFβ-induced αSMA expression in lung fibroblasts derived from IPF patients than did MK-0429 and another benchmarking molecule. Taken together, our work has provided further experimental validation of targeting αv integrin for the treatment of fibrotic lung diseases.

## Results

### Increased αv integrin expression in fibrotic lungs

Recent studies suggested pharmacological targeting of αv integrins is beneficial in multiple tissue fibrosis, including lung, liver, and kidney fibrosis ^6^. To explore the potential role for integrin antagonist in the lung, we first examined the expression of αv integrins in normal human lung fibroblasts (NHLF), normal human bronchial epithelial cells (NHBE), small airway epithelial cells (SAEC), bronchial smooth muscle cells (BSMC), pulmonary artery smooth muscle cells (PASMC), and pulmonary artery endothelial cells (PAEC). The cells were treated with or without TGFβ, a master regulator of myofibroblast activation and extracellular matrix deposition ^7^. The total pool of αv integrins were immunoprecipitated from the cell lysates and subsequently blotted for the abundance of each β subunit. The expression αvβ6 integrin was highly restricted to epithelial cells NHBE and SAEC (**Figure 1A**). αvβ6 is a known epithelium-specific integrin and it binds to latent TGFβ in the extracellular matrix, subsequently inducing local activation of the growth factor and promoting fibrosis ^3,20^. Meanwhile, αvβ3 and αvβ5 were more broadly expressed in a variety of lung cell types (**Figure 1A**). αvβ1, the less-known member of the integrin family, was barely detected in lung fibroblasts (NHLF) and appeared more abundant in epithelial cells (NHBE) and smooth muscle cells (BSMC) upon TGFβ treatment (**Figure 1A**), suggesting that αvβ1 is a highly inducible integrin expressed in multiple cell types. This observation is consistent with our previous report in kidney cell types ^19^, pointing to a potential role for αvβ1 integrin beyond fibroblasts.

**Figure 1.**
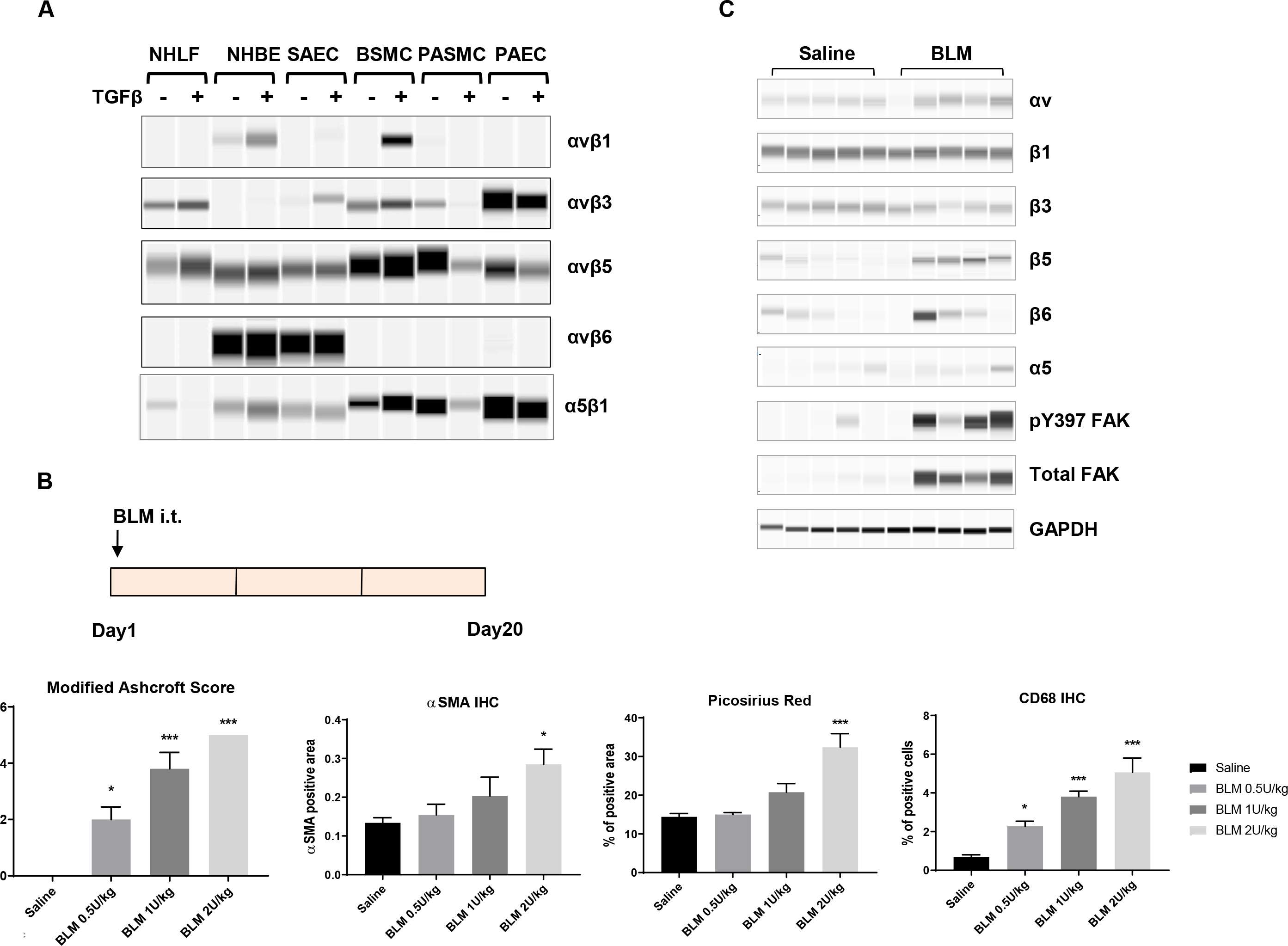
Increased αv integrin expression upon fibrosis induction in the lung. A) The expression of αv integrins in various human primary lung cell types upon TGFβ (5ng/ml for 24 hours) treatment. Following immunoprecipitation with an anti-αv antibody, the αvβ1, αvβ3, αvβ5, and αvβ6 heterodimers were detected by Sally Sue simple western analysis after using antibodies that recognize each individual β-subunit. Normal human lung fibroblast, NHLF; normal human bronchial epithelial cells, NHBE; small airway epithelial cells, SAEC; bronchial smooth muscle cells, BSMC; pulmonary artery smooth muscle cells, PASMC; pulmonary artery endothelial cells, PAEC. Full-length blot images are presented in Supplemental Fig. S6A-E. B) Development of a bleomycin-induced lung fibrosis model in mice. Bleomycin (BLM) was administered at the indicated doses via intra-tracheal (i.t.) instillation. After 20 days, lungs were collected for histological analyses. Modified Ashcroft score, Picosirus red staining, immunohistochemical analyses of αSMA and CD68 of total lung were quantified and shown (mean±SEM, n=5). Immunohistochemistry, IHC. One-way ANOVA followed by Tukey’s test, *p<0.05, **p<0.01, ***p<0.005 vs Saline group. C) integrin expression and signaling in fibrotic lungs (BLM 0.5U/kg bw) was determined by Sally Sue simple western analysis using antibodies that recognized the individual α or β-subunits. GAPDH level in total lung lysates was used as a loading control. Full-length blot images are presented in Supplemental Fig. S6F-N.

Bleomycin-induced lung fibrosis has been commonly used to evaluate the efficacy of a therapeutic agent in preclinical animal studies. After the initial phase of inflammation and cytokine storm, animals develop fibrosis and progressive lung function decline ^21^. To develop the bleomycin model in mice, we administered various doses of bleomycin via intra-tracheal instillation (**Figure 1B**). Twenty days after dosing, lungs were collected for histological evaluation. Bleomycin induced a dose-dependent increase of fibrosis in the lung, as shown by the overall Modified Ashcroft score, αSMA-positive cells, and collagen deposition (Picosirius red staining) (**Figure 1B**). More cells were reactive to CD68 upon bleomycin administration, indicating significant inflammatory macrophage infiltration. Fibrosis and inflammation were evident across all lung lobes evaluated, suggesting wide-spread lesions in the lung (**Supplemental Fig. S1A-B**).

To determine the expression of αv integrins in bleomycin-injured lungs, we next carried out high-throughput Sally Sue simple western analysis of total lung extracts. The abundance of αv, β5, β6, and a5 integrins was significantly upregulated in bleomycin-treated animals, comparing to those from the saline-treated group (**Figure 1C**). Additionally, phospho-Tyr397 and total FAK levels were increased in the bleomycin-treated group, suggesting activation of classical integrin signaling in the lung and providing additional experimental support for αv integrin targeting in this model.

### MK-0429 suppresses fibrosis progression in bleomycin model

We previously found that MK-0429 is an equipotent pan-inhibitor of all αv integrins which reduces proteinuria and renal fibrosis in diabetic nephropathy model ^19^. We next determined whether MK-0429 could inhibit the progression of lung fibrosis *in vivo*. Five days after bleomycin administration, we treated mice with MK-0429 (200mpk via osmotic minipump) or a benchmarking agent (Nintedanib, 60mpk, po, qd). The animals were analyzed for histology and biomarkers after treatment for 14 days (**Figure 2A**). This treatment regimen allowed us to probe the efficacy of the integrin mechanism after bypassing the initial phase of inflammation and cytokine storm.

**Figure 2.**
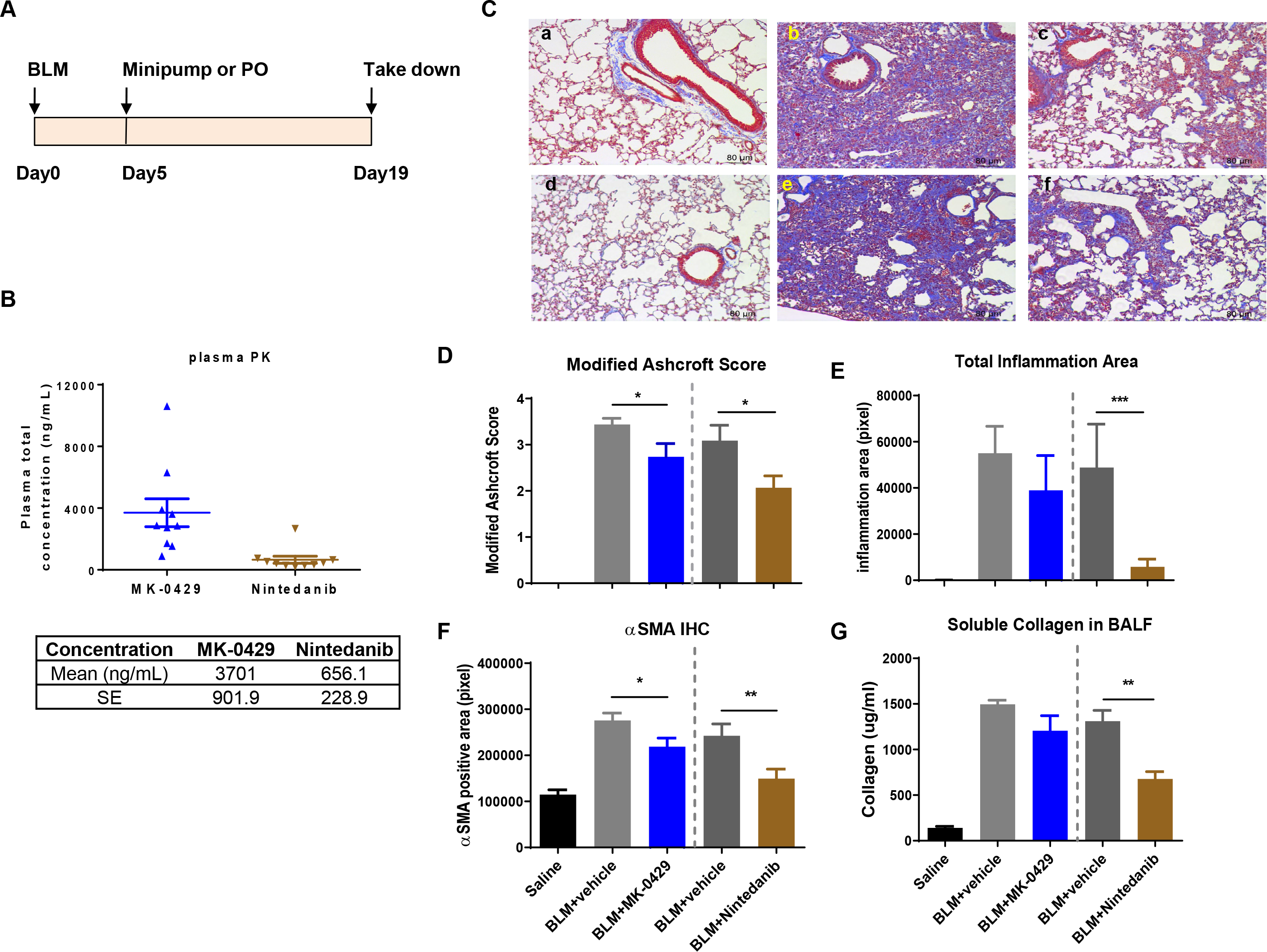
MK-0429 inhibits lung fibrosis in the bleomycin mouse model. A) Schematics of compound administration in BLM model. 5 days after BLM intra-tracheal instillation, the animals were given MK-0429 (200 mpk via osmotic minipump) or Nintedanib (60 mpk po qd). Lungs were collected at Day 19 for histological and biochemical evaluation. B) Plasma total drug concentration was measured 2 hours after final oral dose at Day 19. C) Representative Masson Trichrome staining of mouse lungs. a and d, saline intra-tracheal instillation; b-f, BLM intra-tracheal instillation; a and d, no compound treatment; b, vehicle in minipump; c, MK-0429 in minipump; e, vehicle po; f, Nintedanib po. D) Modified Ashcroft scores of mouse lung. Mean±SEM, n=10. One-way ANOVA followed by Tukey’s test, *p<0.05, **p<0.01, ***p<0.005 vs BLM-vehicle group. E) Total inflammation area in mouse lungs. Mean±SEM, n=10. One-way ANOVA followed by Tukey’s test, *p<0.05, **p<0.01, ***p<0.005 vs BLM-vehicle group. F) Immunohistochemical analysis of αSMA. Mean±SEM, n=10. One-way ANOVA followed by Tukey’s test, *p<0.05, **p<0.01, ***p<0.005 vs BLM-vehicle group. G) Soluble collagen content in bronchoalveolar lavage fluid (BALF). Mean±SEM, n=10. * p<0.05, ** p<0.01, *** p<0.005 vs BLM-vehicle group.

Plasma concentration levels of MK-0429 and Nintedanib were determined at 2 hours after the last dose at the end of the study. The mean plasma concentration was 3701± 902 ng/ml for MK-0429, and 656±229 ng/ml Nintedanib (**Figure 2B**), consistent with the previous report ^19^.

Bleomycin instillation significantly decreased the bodyweight when compared to mice treated with Saline (**Supplemental Fig. S2A-B**). After MK-0429 and Nintedanib administration, there was no significant body weight difference between the bleomycin group and different treatment groups. There was no significant difference for percentage change of body weight among various groups. Mouse lungs were collected at Day 19 for histological evaluation. As shown by the Masson Trichrome staining (**Figure 2C**), control lung tissues had normal lung structure with some collagen around the bronchioles (panels a and d). Administration of bleomycin caused severe multifocal and diffuse fibrosis, thickening of alveolar septa, intra-alveolar fibrosis, and increased perivascular and peribronchiolar infiltration of inflammatory cells (**Figure 2C, panel b**). Nintedanib significantly improved modified Ashcroft score and decreased inflammation after 14 days treatment (**Figure 2C, panel c**; **Figure 2D-E**). Meanwhile, MK-0429 at 200mpk significantly decreased the modified Ashcroft score and had a nonstatistical significant decrease of inflammation in the lung (**Figure 2D-E**). Myofibroblast proliferation in lung tissue was detected by αSMA IHC staining. As shown in **Figure 2F**, bleomycin significantly increased the immunoreactivity for αSMA, both Nintedanib and MK-0429 significantly decreased bleomycin-induced αSMA expression.

Bronchioalveolar lavage fluid (BALF) was also collected for biomarker analyses. Bleomycin increased soluble collagen content and TIMP-1 levels in BALF (**Figure 2G**, **Supplemental Fig. S2C**). Nintedanib significantly decreased BALF soluble collagen and TIMP-1 content after 14 days treatment. The decreases in BALF soluble collagen and TIMP-1 content upon MK-0429 treatment did not reach statistical significance. Together, our results demonstrate that MK-0429 is effective at reducing fibrosis progression in a bleomycin lung injury model.

### Discovery of novel αv integrin monoclonal antibodies with human and mouse cross-reactivity

To obtain new integrin inhibitors for fibrosis, we utilized Adimab yeast surface display platform to screen for tool molecules with better potency than MK-0429. A full-length human naïve IgG library was used for the identification and counter selection of αv-integrin antibodies (**Figure 3A**). Extracellular domains of recombinant human and mouse αvβ1, αvβ3, αvβ5, αvβ6, αvβ8, and α5β1 integrin proteins (**Supplemental Table S1**) were purified, biotinylated, and used as baits to screen for specific binders through several rounds of enrichment of magnetic bead isolation or fluorescence activated cell sorting (FACS). In round 1 and 2, IgG-presenting yeast clones were enriched for target binding and affinity using bait αvβx proteins at a concentration of 50 nM and 500 pM, respectively (**Figure 3B**). In round 3, the best-binders were depleted for α5β1 binding and PSR (poly-specificity reagent) non-specific binding. From the initial screen, we identified 188 unique binders for IgG expression, purification, and functional characterization.

**Figure 3.**
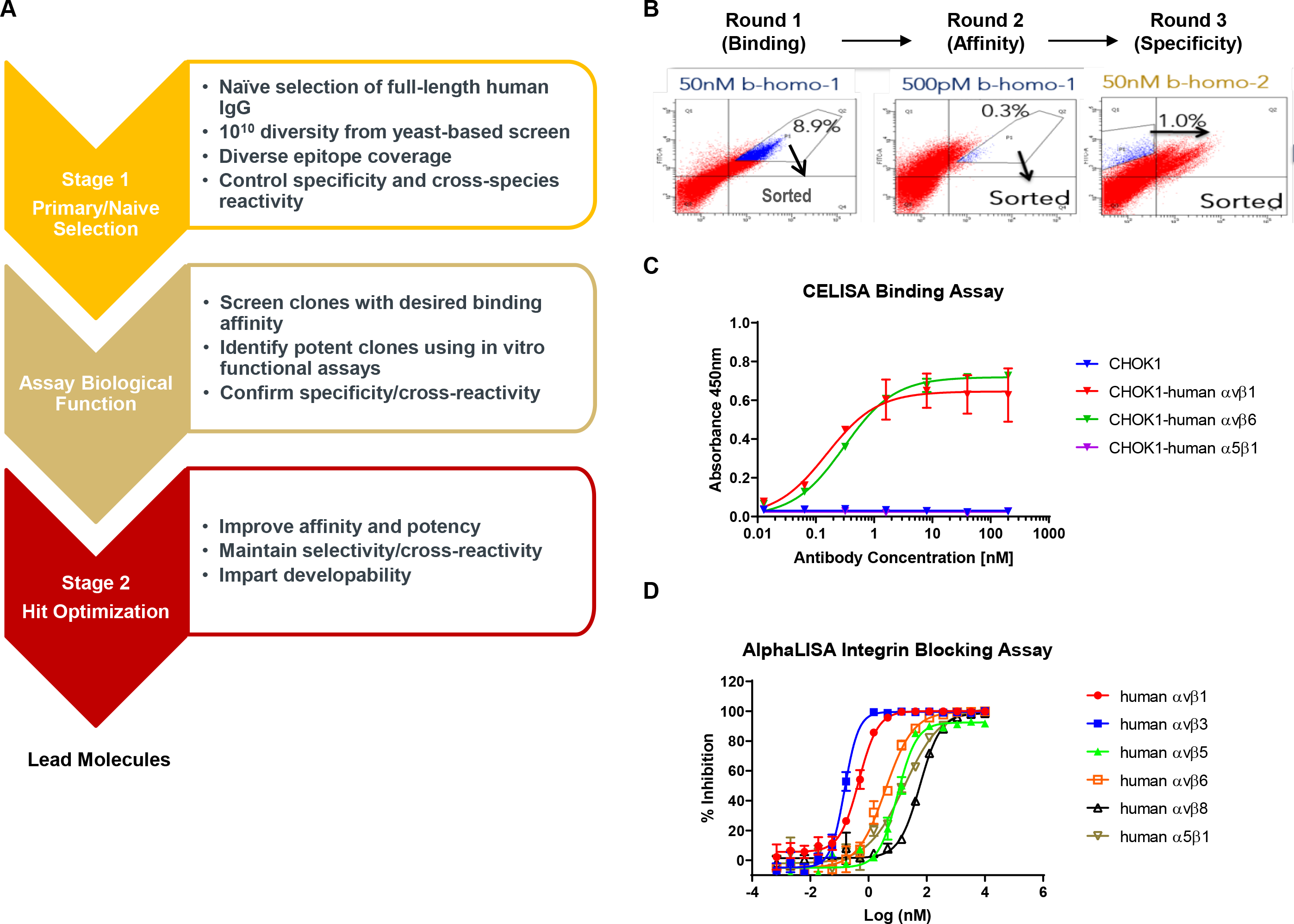
Integrin antibody screening and assay development. 1) Staged efforts to screen integrin antibodies from human naïve IgG library by using Adimab’s yeast display platform. B) IgG-expressing yeast clone selection process. C) Cell-based ELISA (CELISA) binding assays were used as the primary screen for integrin antibody selection. Dose-dependent binding of the benchmarking integrin antibody mAb-24 to various integrin-expressing CHOK1 cells was shown. D) AlphaLISA integrin-ligand binding assays were used for in vitro functional screen. Dose-dependent inhibition of human integrin-ligand binding by MK-0429 in AlphaLISA assay panel.

To examine the binding affinity of each clone, we next generated a set of CHOK1 cell lines that stably expressed human or mouse αvβ1, αvβ3, αvβ5, αvβ6, αvβ8, and α5β1 integrins. The relative abundance of each integrin was determined by FACS analyses after staining the cells with corresponding integrin antibodies (**Supplemental Fig. S3, Supplemental Table S2**). The binding affinity and specificity of each yeast clone to CHOK1 parental cells or CHOK1-αvβ1, αvβ6, or α5β1-expressing cells were determined upon antibody titration in a cell-based ELISA (CELISA) assay. As shown in **Figure 3C**, a benchmarking monoclonal antibody mAb-24 preferentially bound to human αvβ1 and αvβ6 integrin but not α5β1 integrin in this high-throughput cell-based binding assay. 34 antibody clones with 10-fold higher binding affinity towards αvβ1 and αvβ6 integrins were selected for further functional characterization.

We previously determined the in vitro potency and selectivity of MK-0429 by using solid-phase ELISA and thermal shift assays ^19^. To develop a more sensitive and high-throughput functional assay to screen integrin antibodies, we utilized the AlphaLISA platform to develop assays to determine the blocking of integrins-ligand binding by antibodies. It has been shown that fibronectin, vitronectin, and TGFβ latency-associated peptide (LAP) function as endogenous ligands for αvβ1, αvβ3/5, and αvβ6/8 ^5,22–24^. Additionally, α5β1 is a well-known fibronectin receptor ^25^. Thus, we used different ligands for each integrin (**Supplemental Table S3**) and optimized the assay conditions to obtain curve-fitted IC50 values for MK-0429. As shown by the results in **Figure 3D**, MK-0429 demonstrated potent inhibition against human αvβ1 (IC50=1.0±0.3 nM), αvβ3 (IC50=1.6±0.6 nM), αvβ5 (IC50=0.6±0.2 nM), αvβ6 (IC50=2.9±0.6 nM), αvβ8 (IC50=3.1±1.9 nM), and α5β1 (IC50=26.5±5.7 nM), consistent with our previous solid-phase ELISA assay results ^19^. Subsequently, we selected strong binders to examine their abilities to block integrin function in both human and mouse AlphaLISA assays.

Our initial screen led to the identification of 5 unique antibody clones from different germlines that potently inhibited multiple αv-integrins. The top clones were selected for light chain shuffling and affinity maturation on heavy chain CDR1 and CDR2 sequences. Daughter clones with improved binding affinity and potency were expressed in mammalian cells and purified for further characterization to identify lead molecules (**Figure 3A**).

After an initial screening and two rounds of affinity maturation, we identified 5 unique antibodies, Ab-29, Ab-30, Ab-31, Ab-32, and Ab-33, which strongly bound to human αvβ1 and αvβ6 integrins, and in a lesser extent towards α5β1 integrin (**Figure 4A**). Their EC50s of cell-based binding towards each integrin are shown in **Figure 4B**, suggesting that they are pan-αv inhibitors. Notably, those five molecules were also reactive to mouse αvβ1, αvβ6, and α5β1 integrins (**Figure 4B**, **Supplemental Fig. S4A**). To date, there are no αv-integrin antibodies with human and mouse cross-reactivity reported. We next carried out AlphaLISA integrin blocking assays to examine the effect of each antibody on integrin-ligand binding. As shown in **Figure 4B**, Ab-29, Ab-30, Ab-31, Ab-32, and Ab-33 substantially inhibited human αvβ1, αvβ3, αvβ5, αvβ6, αvβ8, and α5β1 integrins, with IC50s comparable to those of MK-0429 and the benchmarking mAb-24. MK-0429 was potent at blocking mouse integrin-ligand binding in all cases except mouse αvβ6 and α5β1, where it showed weaker activities (**Figure 4B**, **Supplemental Fig. S4B**). Ab-29, Ab-30, Ab-31, Ab-32, and Ab-33 also strongly inhibited mouse αvβ1, αvβ3, αvβ6, and αvβ8 integrins. Their activities towards mouse α5β1 integrin were negligible. The binding affinity (Kd) of each antibody to selected integrin proteins was within the range of 2-48 nM range as assessed by the ForteBio Octet Red system (**Supplemental Fig. S4C**). Epitope binning revealed unique binding sites of our antibodies compared to other known αv-integrin antibodies (data not shown). Overall, our molecules represent a new class of αv-integrin blocking antibodies that could be used as mouse surrogates for rodent preclinical studies.

**Figure 4.**
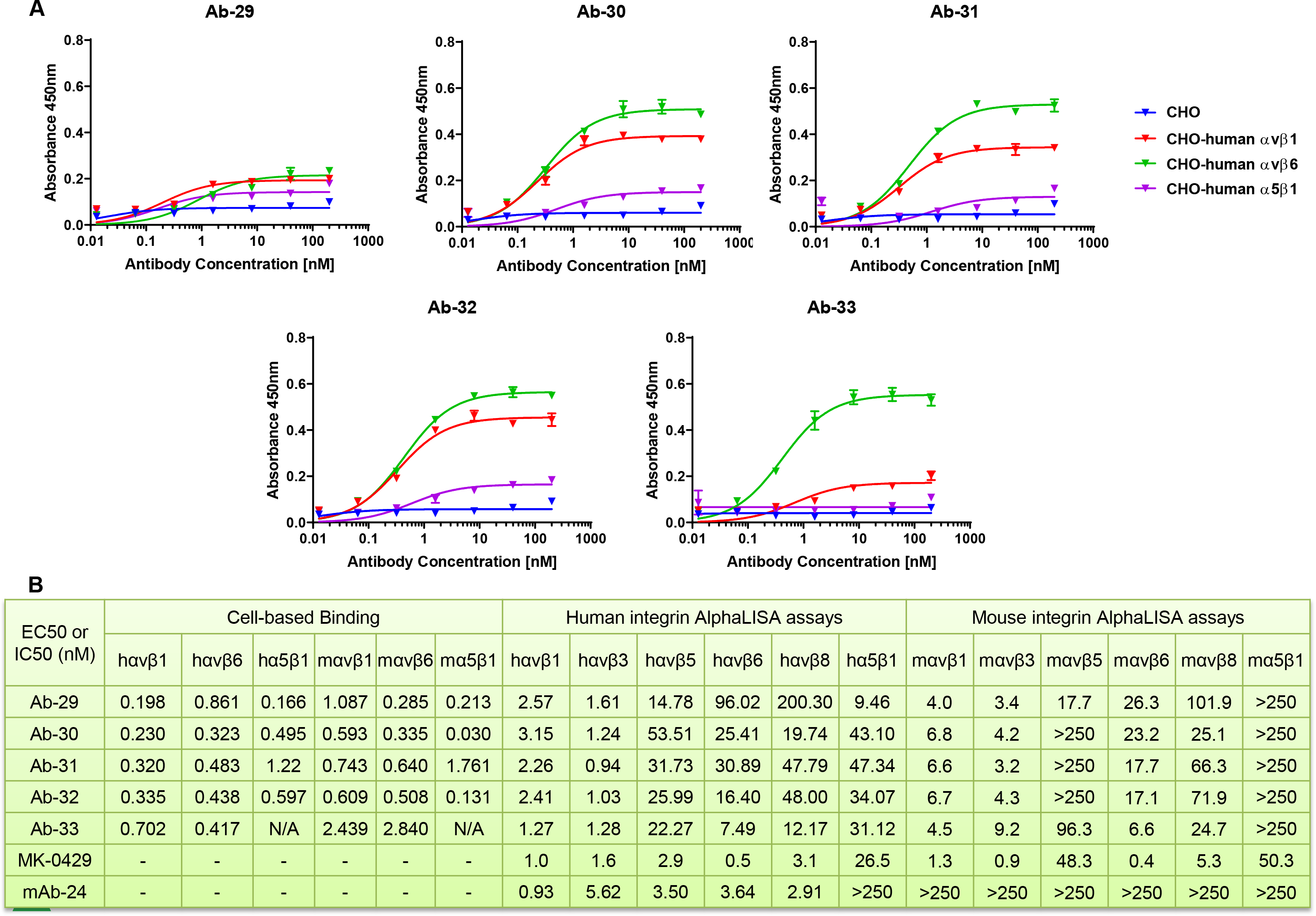
Discovery a set of antibodies with strong blocking activities against both human and mouse αv integrins. A) Titration of Ab-29, Ab-30, Ab-31, Ab-32, and Ab-33 for their binding to CHOK1-human αvβ1, αvβ6, and α5β1 stable cell lines in CELISA assays. B) EC50 of Ab-29, Ab-30, Ab-31, Ab-32, and Ab-33 in CELISA assays, as well as their IC50 in human and mouse AlphaLISA integrin blocking assays.

### Ab-31 blocks integrin-mediated cell adhesion

Previous studies utilized cell adhesion assays to determine integrin subtype selectivity and cellular function ^5,26^. This cell-based assay measures the binding of ligand to integrins that are endogenously present or over-expressed on the cell surface. Compared to in vitro functional assays using recombinant protein, cell adhesion results are more reflective of native integrin-ligand binding conformation. We generated CHOK1 stable cell lines that expressed various human and mouse αvβx integrins (**Supplemental Fig. S3**). Both αvβ1 and α5β1 can function as the fibronectin receptor in cell adhesion assay ^25,27^. To delineate the role of αvβ1 in cell adhesion, we deleted endogenous hamster α5 gene in CHOK1 cells via CRISPR knockout/KO technology, and subsequently overexpressed αvβ1 to generate a CHOK1-α5KO-αvβ1 stable line. We next examined the effect of MK-0429 and our top monoclonal antibody clone, Ab-31, on the adhesion of cells to fibronectin or vitronectin matrix. MK-0429 showed little inhibitory activity in CHOK1 parental cells on fibronectin; however, it potently reduced the adhesion of mouse αvβ1-expressing CHOK1-α5KO cells (**Supplemental Fig. S5A**). Cell adhesion of mouse αvβ3 and αvβ5 on a vitronectin matrix were also decreased upon MK-0429 treatment. Similarly, Ab-31 significantly inhibited mouse αvβ1, αvβ3, and αvβ5 integrin-mediated cell adhesion, with IC50s of 1.5 nM, 1.0 nM, and 5.6 nM respectively (**Figure 5A**). The cell-based assays showed that Ab-31 is a potent αv-integrin inhibitor in a setting that resembles native integrin conformation.

**Figure 5.**
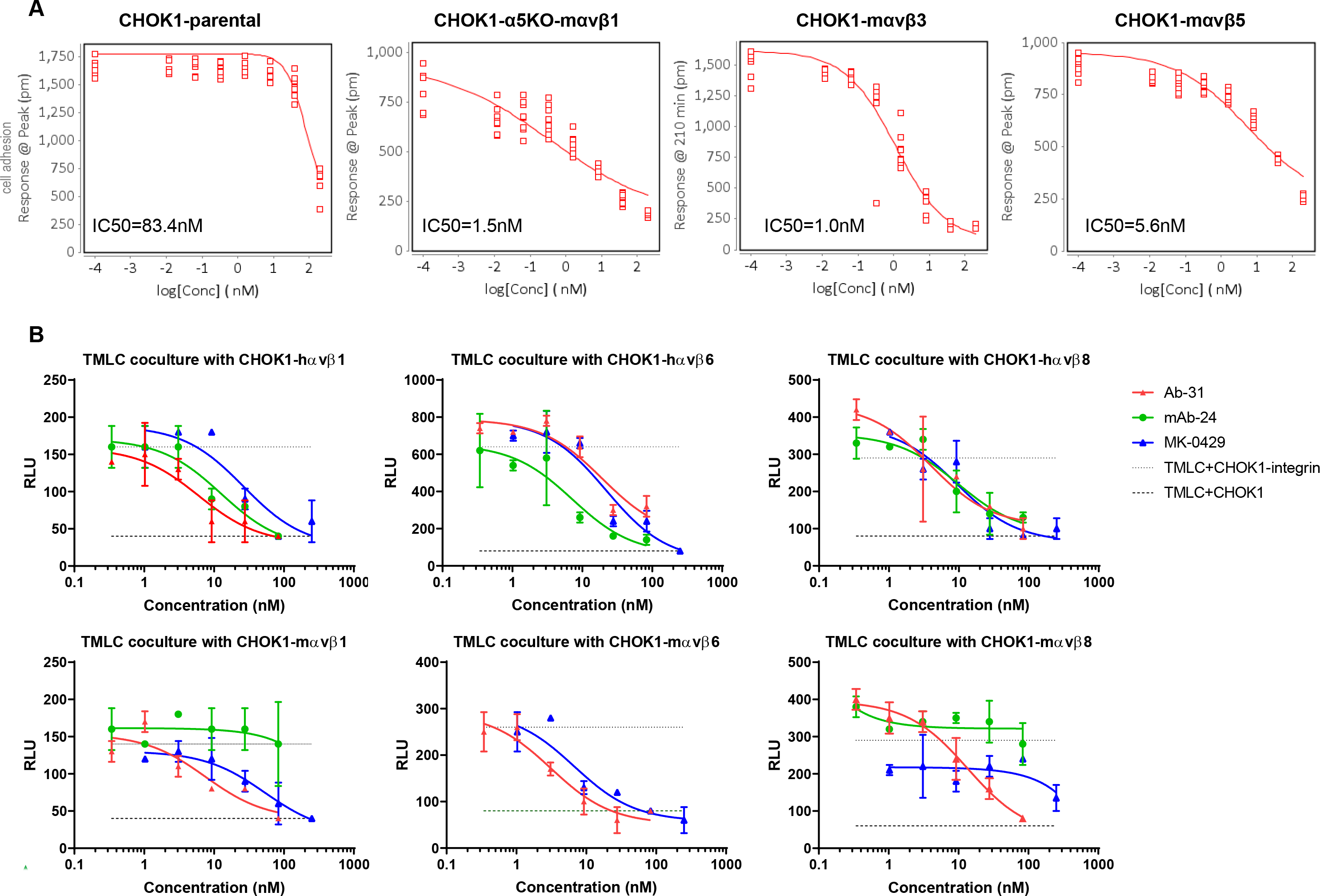
Ab-31 inhibits integrin-mediated cell adhesion and latent TGFβ activation. A) The effect of Ab-31 on the adhesion of CHOK1 parental, CHOK1-α5KO-mαvβ1, CHOK1-mαvβ3, and CHOK1-mαvβ5 cells to fibronectin or vitronectin matrix. B) pan-αv integrin inhibitors suppress latent TGFβ activation in the transfected Mink lung epithelial cells (TMLC) and CHOK1-integrin co-culture system. The effects of Ab-31, MK-0429, and benchmarking mAb-24 on PAI-1 luciferase activity were shown.

### Ab-31 substantially inhibits integrin-mediated latent TGFβ activation

The epithelium-specific αvβ6 integrin blocks the activation of latent TGFβ ^3,8^, a major mechanism-of-action for αv-integrin to modulate fibrosis. Rifkin et al. transfected mink lung epithelial cells with a firefly luciferase reporter under the control of PAI-1 (plasminogen activator inhibitor-1) promoter ^28^. The transcription of PAI-1 is tightly controlled by the TGFβ-Smad pathway ^29,30^. When co-culturing these transfected mink lung epithelial cells (TMLC) with another cell type-expressing integrin, the luciferase activity is driven by the abundance of mature TGFβ presented in extracellular matrix and culture medium. This co-culturing system provides a sensitive measurement of latent TGFβ activation by integrins.

In addition to αvβ6 integrin, both αvβ1 and αvβ8 have been shown activating latent TGFβ ^5,11,31^. We next examined the activity of our monoclonal antibody Ab-31 against TGFβ activation in the TMLC-integrin co-culture system. Notably, Ab-31 substantially blocked the activation of latent TGFβ by human and mouse αvβ1, αvβ6, and αvβ8 integrins, with IC50 at 6.1 nM, 19.1 nM, 3.9 nM, 7.5 nM, 3.1 nM, and 13.5 nM, respectively (**Figure 5B**). MK-0249 also demonstrated strong inhibitory effect at TGFβ activation except in a mouse αvβ8 co-culture assay. The benchmarking antibody, mAb-24, was potent in human αvβ1, αvβ6, and αvβ8 co-culture systems, but elicited minimal activity against mouse integrins, which is consistent with its activities in AlphaLISA assays. Together, our results found that pan-αv integrin monoclonal antibody Ab-31 strongly inhibits integrin-mediated cellular functions, including cell adhesion and latent TGFβ activation.

### Ab-31 demonstrates superior inhibitory activity against TGFβ-induced αSMA expression

In addition to the inhibition of latent TGFβ activation, αv-integrin inhibitors also function downstream of TGFβ signaling ^19^. To further characterize the activities of our molecules in a fibrosis-relevant cell type, we next examined the expression of αSMA in primary human lung fibroblasts. Normal human lung fibroblasts were cultured, treated with TGFβ, and stained for αSMA by high-content phenotypic imaging analysis (**Figure 6A**). Upon TGFβ induction, the fluorescence intensity of αSMA was vastly increased (**Figure 6A**), and αSMA-associated cellular morphological changes were recognized as consistent with cell stimulated by the machine learning-based STAR software (Harmony high-content imaging and analysis software, PerkinElmer Inc.). Pretreating cells with MK-0429or with SB-525334, a potent inhibitor of TGFβ type I receptor ALK5 (activin receptor-like kinase) ^32^ significantly reduced αSMA intensity and the percentage of stimulated cells following TGFβ treatment.

**Figure 6.**
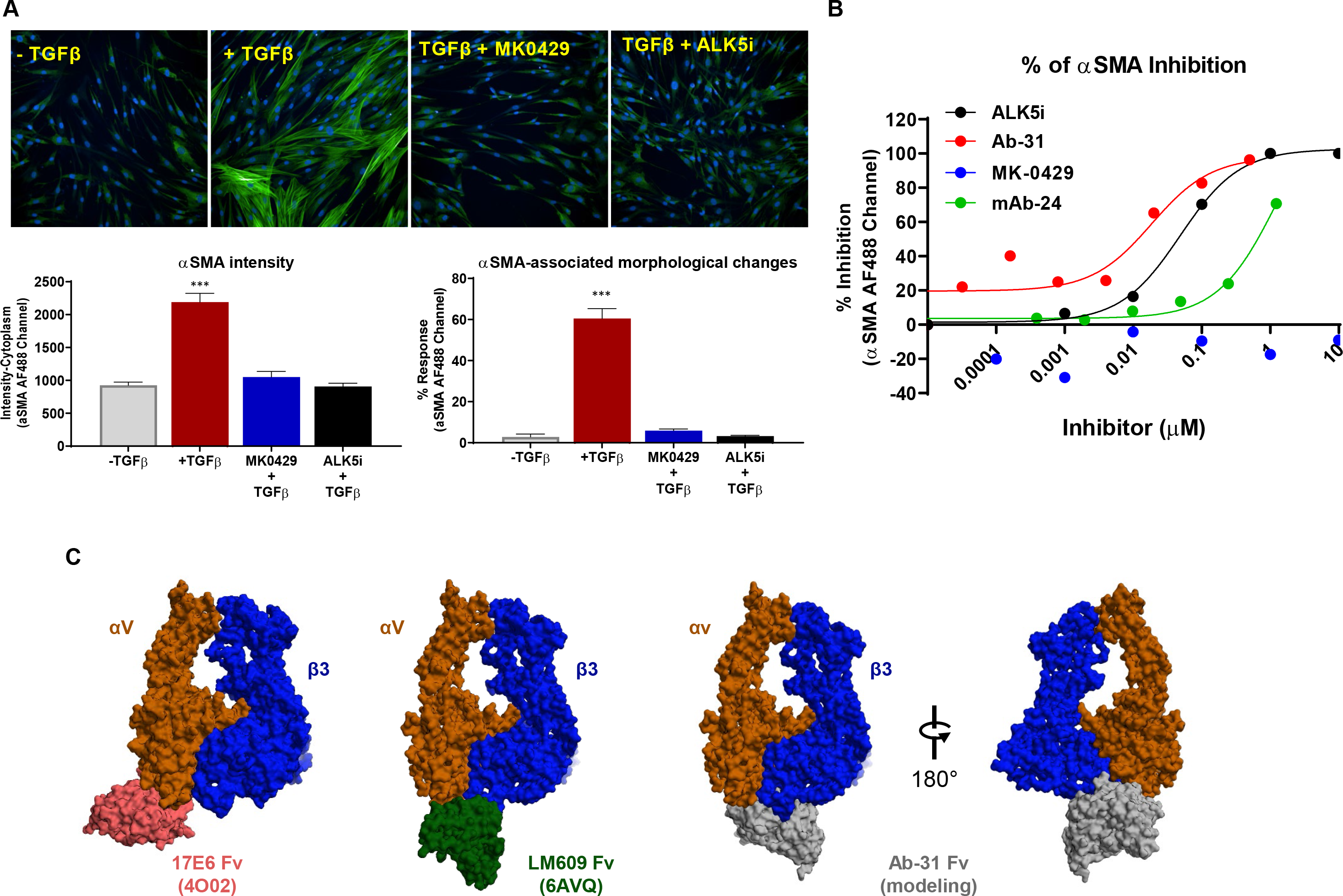
Ab-31 reduces TGFβ-induced αSMA expression in lung fibroblasts. A) Normal human lung fibroblasts were stimulated with TGFβ (5 ng/ml) and stained with anti-αSMA antibody for immunofluorescence analysis. Cells were pre-treated with or without MK-0429 (10 μM) or the ALK5 inhibitor SB-525334 (10 μM) 30 minutes before the addition of TGFβ. 48 hours after the treatment, cells were imaged with Opera Phenix high-content screening system for αSMA expression (fluorescence intensity) and αSMA-associated morphological changes (STAR program). B) The effects of Ab-31, MK-0429, and benchmarking antibody mAb-24 on TGFβ-associated αSMA induction in IPF patient lung fibroblasts. C) Structural modeling predicts a distinct integrin binding mode for Ab-31. The structures of two therapeutic monoclonal antibodies, Abituzumab (17E6, RCSB PDB: 4O02) and LM609 (RCSB PDB: 6AVQ), in complex with αvβ3 integrin were shown to highlight the difference in binding modes for each molecule. A model of Ab-31 and αvβ3 integrin complex was determined by docking of related antibody sequence to αvβ3 structure (see Methods). For visualization, only the Fv region of each antibody were shown.

After establishing this αSMA phenotypic assay platform, we next examined the effects of integrin inhibitors on lung fibroblasts derived from IPF patients. As seen in the healthy human fibroblast assays, the expression of αSMA and percentage of stimulated cells among fibroblasts from IPF patients were also increased after TGFβ treatment (**Figure 6B**). The ALK5 inhibitor, SB-525334, potently inhibited TGFβ-mediated αSMA expression in patient fibroblasts, with IC50 of 56±22 nM. Notably, MK-0429 had minimal activity at αSMA induction even at a concentration of 10 μM, suggesting that the network for αSMA regulation is distinct in IPF patient fibroblasts than that of normal human lung fibroblasts. The benchmarking monoclonal antibody mAb-24 was also less effective at inhibiting αSMA expression with IC50 above 1 μM. Interestingly, Ab-31 demonstrated a strong dose-dependent inhibition of αSMA intensity and associated morphological changes with IC50 of 33±21 nM, similar to that of the ALK5 inhibitor (**Figure 6B**). It is noteworthy that Ab-31 was ineffective in normal human lung fibroblasts (data not shown), raising a possibility that this molecule preferentially recognizes integrin confirmation under fibrotic state. Our data demonstrate that Ab-31 was a potent inhibitor of TGFβ signaling in IPF patient lung fibroblasts and exhibits superior anti-fibrotic activity than MK-0429 or other comparator in cell-based assays.

### Structural modeling of integrin-antibody complex

Over the past 20 years, a diverse set of integrin monoclonal antibodies have been discovered and extensively characterized that can elicit inhibitory, stimulatory, or neutral activity upon integrin binding, ^33^. Integrin antibodies are also known to recognize active conformational epitope, such as 12G10 and 9EG7 clones of β1-integrin ^34–36^. To further understand the mechanism of action of our antibody, we next set out to predict how it interacts with integrins.

The first crystal structure of integrin heterodimer was solved in 2001 and revealed a bent inactive confirmation of αvβ3 integrin ectodomains ^37^. Since then, several integrin-antibody complex structures have been determined and have provided valuable information in regard to their regulatory function. 17E6 (Abituzumab) is a therapeutic pan-αv integrin antibody that has been tested in multiple Phase 2 clinical trials in cancer patients and systemic sclerosis patients with interstitial lung disease ^38^ (clinicaltrials.gov identifier NCT02745145). The crystal structure of the 17E6 Fab fragment in complex with αvβ3 revealed that the antibody exclusively bound to the αv subunit and helped to establish a potential allosteric inhibition mechanism via steric hindrance (PDB: 4O02) ^39^. As illustrated in the left panel of **Figure 6C**, the variable region (Fv) of 17E6 binds to an epitope outside the ligand binding interface of the α- and β-subunits. In contrast, LM609 is an αvβ3-specific blocking antibody of which the humanized variants, Vitaxin and Etaracizumab (Abegrin), have been tested in several clinical trials for oncology indications ^40–42^. A combination of crystallographic and cryo-EM work from Veesler et al. found that LM609 bound to the apex of the integrin headpiece close to but without directly obstructing the RGD-binding site ^43^. The structure of LM609 Fv fragment in complex with αvβ3 integrin is illustrated in the second left panel of **Figure 6C**, with an epitope locating at the interface between the α- and β-subunits. Compared to either of these characterized antibodies, Ab-31, as well as other top clones from our screen, represented a unique class of pan-αv integrin antibodies with human and mouse cross-reactivity. To better understand the mechanism of integrin inhibition by Ab-31, we set out to model the Ab-31-integrin complex through homology modeling and docking. Using the antibody Fv sequence, we generated a series of 10 models for the Fv structure of Ab-31 using MOE (Chemical Computing Group). The lowest energy structure was then docked against the known structure of the known αvβ3 structure (see Methods). Following visualization and filtering of potential poses against known experimental variants, the final pose shown in **Figure 6C** was determined. Interestingly, the model predicted that Ab-31 bound directly at the interface between α- and β-subunits and somewhat blocked access to the RGD-ligand binding site (**Figure 6C**, right panels), indicating a distinct mode of action compared to 17E6 and LM609. This model is consistent with our epitope binning findings that Ab-31 had an epitope that did not overlap with other integrin antibodies (data not shown). Further structural and biophysical characterization will help to better understand its mode of action and cross reactivity.

## Discussion

Fibrosis, characterized as excessive accumulation of extracellular matrix in the parenchyma, is a hallmark of end-stage organ failure and is strongly associated with poor outcome in patients with heart failure and chronic kidney diseases ^44–46^. A therapeutic agent targeting fibrosis could potentially delay disease progression and offer synergetic benefits when combined with standard-of-care agents. In 2014, FDA approved the use of Pirfenidone and Nintedanib for lung fibrosis. Both Pirfenidone and Nintedanib slow down lung function decline; however, neither drug was able to offer symptom relief (breathing difficulty) or a substantial survival benefit ^47^. There is an unmet medical need to develop new IPF therapies that bring clinically meaningful efficacy to patients.

IPF is multi-factorial disease and the dominant mechanism that drives IPF pathogenesis is unclear. The mechanism of action for Pirfenidone is presently unknown, likely involving multiple pathways that include anti-inflammation and TGFβ suppression ^48^. Nintedanib is an inhibitor for multiple receptor tyrosine kinases, such as VEGFR, PDGFR, and FGFR ^49^. Currently, there are several mechanisms being tested in the clinic for IPF patients, including but not limited to the CTGF antibody Pamrevlumab, the Autotaxin inhibitor GLPG-1690, and recombinant Pentraxin 2 (PRM-151). In recent years, integrin inhibitors have emerged as key mediators of tissue fibrosis. In particular, αv-containing integrins, such as αvβ6, modulate local TGFβ activation and myofibroblast activation with strong preclinical validation for lung fibrosis ^3,9,50^. In the present study, we used MK-0429 as a tool molecule to further validate the role of αv integrins in preclinical lung fibrosis model. Furthermore, we initiated an antibody discovery campaign and discovered a set of novel integrin monoclonal antibodies with human and mouse cross-reactivity. Among these, Ab-31 potently blocked integrin-ligand binding, inhibited integrin-mediated cell adhesion, suppressed both the activation of latent TGFβ and the αSMA expression induced by activated TGFβ. Notably, Ab-31 demonstrated superior activity at inhibiting TGFβ response in IPF patient lung fibroblasts compared to MK-0429 and another comparator molecule, making it a unique tool molecule to study anti-fibrotic efficacy in vivo. It is intriguing that Ab-31 and MK-0429 have comparable activities when tested using in vitro AlphaLISA integrin-ligand blocking assays, but their impact on TGFβ signaling in IPF patient lung fibroblasts was drastically different. In previous reports, bivalent 17E6 and LM609 were postulated to interfere with integrin clustering and internalization on the cell surface, enhancing their therapeutic effects over that of a monovalent Fab fragment ^39,43^. It is possible that Ab-31 also functions by regulating integrin clustering and internalization. Further structural and biophysical characterization is warranted to elucidate the inhibitory mechanism of Ab-31.

The complexity of integrin biology and the overlapping roles of multiple integrins in the progression of the disease suggest that a pharmacological pan inhibitor would be beneficial, leading to clinically meaningful inhibition of fibrosis. Interestingly, a recent GWAS study of large population provides strong genetic evidence that supports targeting the αv integrin to improve lung function, potentially expanding the use of αv integrins inhibitors to a broader patient population, such as chronic obstructive pulmonary disease (COPD) ^13^. The poly-pharmacology nature of an αv inhibitor raises potential safety concerns. Notably, MK-0429 has been tested in the clinic in osteoporosis patients over the duration of 52 weeks with relatively well-tolerated safety profile ^16,51^. One main mechanism-of-action of integrin inhibitors is to inhibit latent TGFβ activation. Compared to a cardiovascular safety signal of TGFβ inhibitor, the safety profile of pan-αv integrin inhibition is more tolerable. Recently, an αvβ6 antibody (BG00011) was withdrawn from phase 2 clinical trials in IPF patients due to safety issues (clinicaltrials.gov identifier NCT03573505). It will be of interest to compare the efficacy and safety profiles of αvβ6-specific and pan-αv inhibitors. The preferential effect of Ab-31 on IPF patient lung fibroblasts over normal human lung fibroblasts suggested that this antibody may selectively recognize either an active or diseased-associated integrin conformational state, making it a promising antagonist with an improved therapeutic index.

In summary, our present work describes the discovery a new class of integrin monoclonal antibodies that potently inhibit integrin-ligand binding, integrin-mediated cell adhesion, and TGFβ signaling. Our molecules exhibit distinct human and mouse cross-reactivity and structural modeling predicts a unique mode of inhibition. Future computer-aided rational design will allow the development of optimized molecules for further anti-fibrotic therapy.

## Acknowledgements

The authors thank Zubia Naji for project management support and thank Xiaolan Shen, Audrey Chu, Dana Nojima, Mohammad Tabrizifard, Maarten Hoek, John Hunter, Ednan Bajwa, Paul Coleman, and Gordon Wollenberg for their valuable feedback throughout the work. We thank Dr. Jesse Nussbaum for his critical review of the manuscript. We thank Drs. Roy Zent and Ambra Pozzi at the Vanderbilt University for their valuable feedback. We thank Dr. Daniel Rifkin at New York University for his expertise and providing mink lung epithelial cells for latent TGFβ assays. Histological staining was provided by HistoBridge via Science Exchange. We acknowledge the dedicated assay development work of the late Mrs. Kim O’Neill while fighting late-stage cancer.

## Author contributions

J.Z., M.G-C., J.H., T-Q.C., T.A., S.P., J.C.M., Z.R., H.H.S., and M.H. conceived the experiments; J.Z., T.W., A.S., J.J., J.M., S.T., M.J.E., W.M., K.O., D.G.L, T.H., Q.Z., and W.D. performed experiments; E.C-J. and T-Q.C. coordinated in vivo studies; S.A.H. and A.C.C. performed antibody structural modeling, J.Z., A.S., J.J., J.M., S.T., S.A.H., M.J.E., E.C-J., H.Y.M., A.C.C., J.C.M., H.H.S., and M.H. wrote the manuscript; T.G., and S.T. provided critical expertise and feedback.

## Competing interests

The authors are/were employees of Merck Sharp & Dohme Corp., a subsidiary of Merck & Co., Inc., Kenilworth, NJ, USA and/or shareholders of Merck & Co., Inc., Kenilworth, NJ, USA.

## Data Availability

All data generated or analyzed during this study are included in this manuscript and its supplementary information files.

## Methods

### Cultured Cells and Reagents

Normal human lung fibroblasts (Lonza) and Primary Human IPF Lung Parenchymal Fibroblasts (Donor2) (BioIVT, #PCR-70-0214) were incubated at 37°C and 5% CO2. To induce fibrotic response in primary cells, a final concentration of 5ng/ml of recombinant human TGFβ1 (BioLegend, #580702) was added to the culture media and treated for 24~48 hours. Anti-TGFβ neutralizing antibody (1D11) and the isotype control mouse IgG1 were from BioXcell. SB-525334 and bleomycin were from Sigma. MK-0429 was synthesized by Merck & Co., Inc., Kenilworth, NJ, USA. CHOK1-integrin stable lines were cultured in DMEM/F12, Glutamax (Gibco #10565018), 10% FBS (Gibco 310091148), 1x Pen/Strep (Gibco 315140-148), and 6 ug/mL Puromycin (Gibco #A1113803).

### Recombinant integrin proteins

The full extracellular domains for all α and β integrin subunits (human and mouse versions, **Supplemental Table S1**) were codon optimized for mammalian expression (Genewiz, NJ) and inserted into the HindIII and XhoI sites of pcDNA3.1. αv and α5 contained a C-terminal (GGGS)_3_ linker with an acidic coiled-coil with a cysteine for disulfide-bond formation, a GG-Avitag (Avidity, CO), and a hexahistidine tag. β1, β3, β5, β6, and β8 contained a C-terminal (GGGS)_3_ linker with a basic coiled-coil with a cysteine, and a GG-Avitag. For expression, 10 L of Expi293 cells (ThermoFisher, standard protocol) were co-transfected with 0.5 mg/L each of both an α and a β subunit, either human (h) or mouse (m) and grown for 72 hours at 37°C in shake flasks (hαvβ1, hαvβ3, hαvβ5, hαvβ6, hαvβ8, hα5β1, mαvβ1, mαvβ3, mαvβ5, mαvβ6, mαvβ8, mα5β1). Media was harvested by centrifugation and soluble supernatant concentrated to 1 L by TFF (tangential flow filtration, Pellicon 10K) in 25 mM Tris pH 8.0, 300 mM NaCl, 40 mM Imidazole. Sample was centrifuged again (15 minutes @ 3500 g) prior to purification. For expression, the sample was loaded over a HisTrap FF (2 x 5 mL) column pre-equilibrated in 25 mM Tris pH 8.0, 300 mM NaCl, 40 mM Imidazole, washed for 15 CV with the equilibration buffer, and eluted with a gradient from 40-500 mM Imidazole (in 25 mM Tris pH 8.0, 300 mM NaCl). Protein not needing biotinylation was further purified using a Superdex 200 column in 25 mM TRIS pH 8.0, 150 mM NaCl, 1 mM MgCl_2_, 1 mM CaCl_2_. Eluted fractions were concentrated to 2 mg/mL, aliquoted and frozen. For biotinylation, sample that needed to be biotinylated after elution from the initial HisTrap FF column was concentrated to 3.0 mg/mL and desalted/buffer exchanged (ZebaSpin desalting column) into 25 mM Tris pH 8.0, 150 mM NaCl. Sample was combined with Biomix B and BirA (as per Avidity protocol) and incubated at 4°C for 14 hours. Biotinylated sample was further purified using a Superdex 200 column in 25 mM TRIS pH 8.0, 150 mM NaCl, 1 mM MgCl_2_, 1 mM CaCl_2_. Eluted fractions were concentrated to 2 mg/mL, aliquoted and frozen. Biotinylation was verified by either streptavidin binding gel shift, or with the Pierce Fluorescence Biotin Quantification Kit (#46610).

### Antibody Discovery, Optimization, and Production

*De novo* antibody discovery for αv-integrins were executed on pre-immune Adimab yeast display libraries with a diversity of 10^10 ^52^. The soluble proteins used in the yeast display selections are biotinylated recombinant integrin ectodomain proteins described in **Supplemental Table S1**. All proteins were analytically and biophysically verified by binding against known integrin antibodies (**Supplemental Table S2**), SEC, SDS-PAGE, and endotoxin. Briefly, a yeast IgG library was subjected to multiple rounds of selection by magnetic activated cell sorting (MACS) and florescence activated cell sorting (FACS, BD ARIA III) in PBS buffer containing 1mM MnCl_2_. Selections were performed using 100 nM human or mouse αvβ1 followed by rounds of enrichment of populations that were cross-reactive to 100 nM of αvβ3, αvβ5, αvβ6, and αvβ8. Along with the selection progress, the decreased antigen concentrations are also applied for enhancing selection pressure. Due to the high sequence homology among all integrin isoforms, isoform-specific selections were achieved by negative sorting of α5β1 to collect the population that are not binding to α5β1. Top clones were isolated by affinity maturing its parental clone through shuffling the light chain and optimizing heavy chain CDR1 and CDR2 sequences. The selection of optimization libraries was repeated using 10 nM αvβ1 followed by enrichment of cross-reactivity to αvβ3, αvβ5, αvβ6, and αvβ8, but not α5β1. The isolated clones were then sequenced to identify the unique antibodies and screened for isoform binding profiles by Octet Red. The heavy chain and light chain genes of top clones were cloned into pTT5 mouse Fc-mutated IgG1 backbone vector and produced in Chinese Hamster Ovary (CHO) cells and purified using protein A chromatography. Antibodies were formulated at 2 mg/mL in a buffer composed of 20 mM sodium acetate and 9 % sucrose (pH 5.5). Isotype control antibodies was used in subsequent assays.

### Integrin Cell-based ELISA (CELISA) assay

Cell binding EC50 data for antibodies were obtained for CHOK1 parental cells and CHOK1 cells expressing human or mouse αvβ1, αvβ6, and α5β1 integrins by cell-based ELISA. Three days prior to the assay cells were seeded at 10,000 cells per well in 100 μL/well of media in Falcon 96-well Flat-Bottom Tissue Culture Plates (REF 353916), resulting in an even cell monolayer in each well on the day of the assay. On the day of the assay, media was removed from the cells and a 5-fold titration of primary antibodies was added at concentrations ranging from 0 to 200 nM in TBSF+MnCl_2_ buffer (25 mM Tris, 0.15 M NaCl, 0.1% BSA, 0.5 mM MnCl_2_, pH = 7.5). Cells incubated with primary antibody solution for 1 hour at room temperature. Primary antibody was removed and plates were washed twice with 150 μL/well of 1x DPBS+Tween20 (TEKnova Cat # P0297, diluted to 1x in distilled water GIBCO #15230-147), before adding 50 μL/well of a 1:5000 dilution of HRP-conjugated goat anti-mouse IgG (Southern Biotech, #1030-05) in TBSF+MnCl_2_. After 30 minutes of incubation at room temperature, secondary antibody was removed, and cells were washed twice with 150 μL/well of 1x DPBS+Tween20. 50 μL/well of TMB substrate (1-Step Ultra TMB-ELISA, Thermo Scientific, #34028) was added, incubated with cells at room temperature and 50 μL/well of TMB stop solution (Seracare, #KPL 50-85-05) was added after 5 minutes. Absorbance was read on a TECAN plate reader at 450 nm, with a 620 nm reference wavelength.

### Integrin AlphaLISA assay

In the AlphaLISA integrin assays, we used a HEPES based buffer (25 mM HEPES pH 7.4, 137 mM NaCl, 1 mM MgCl_2_, 1 mM MnCl_2_, 2 mM CaCl_2_, 2.7 mM KCl, and 0.05% Tween-20). Optimal assay conditions were determined with a titration of the reagents for signal noise ratio (S/N), linear range, etc. For the respective integrins, we used the final concentrations described in **Supplemental Table S3**. Briefly, the testing antibody (8 μL) was added as a 4x (of Final testing concentration) solution in the HEPES buffer described previously into a 384w AlphaPlate (Perkin Elmer #6005350). Then the integrin and ligand were added sequentially as 8x (4 μL each). Plates were sealed and incubated at room temperature for 2 hours. Then 4x acceptor bead solution (8 μL) was added, and the plates were resealed and incubated at RT for 1 hour. Finally, 4x (8 μL) donor beads solution was added in a darkened room and incubated for 45 minutes at room temperature. Plates were read on the Envision (Perkin Elmer) in AlphaScreen mode within 3 hours of donor bead addition.

### Cell adhesion assay

For plate preparation, Epic 384-well assay plates (Corning #5040) were washed with OptiMEM (Gibco #31985-070) and coated with murine fibronectin (Abcam #ab92784) at a concentration of 0.1 ug/well for CHO-a5KO-mαvβ1 cells; murine vitronectin (Abcam #ab92727) at 0.075 ug/well for CHO-mαvβ3 cells; or murine vitronectin at 0.125 ug/well for CHO-mαvβ5 (all at a volume of 25 μl/well in OptiMEM), for 1 hour. Coating solution was then removed by flicking and plates were washed with OptiMEM and blocked with 25 ul/well of assay buffer (1% Ovalbumin (Sigma #A5503-10G) in OptiMEM containing 1X Pen-Strep (Gibco #15140-148)). A 5-minute baseline reading was taken on a Corning Epic Plate Reader (Model: Epic BT-157900) before removing the assay buffer and adding cells pre-incubated with inhibitors. For inhibitor titration preparation: 5-fold serial dilutions of small molecule inhibitor MK-0429 or antibody inhibitor Ab-31 were prepared in assay buffer at 4X the final concentration in a 384-well polypropylene plate in a volume of 15 μl per well, starting at a concentration of 800 nM. For cell preparation: On the day of the assay, frozen cell stocks were thawed and transferred to 10 mL complete media (DMEM/F12 + Glutamax (Gibco #10565-018) containing 10% FBS (Gibco #10091-148) and 1X Pen-Strep (Gibco 315140-148) for CHO and CHO-mαvβ3 cells, or F12K Nutrient Mixture (Gibco #21127-022) containing 10% FBS (Gibco #10091-148) and 1X Pen-Strep (Gibco #15140-148) for CHO-α5KO-mαvβ1 and CHO-mαvβ3 cells at 37°C in a 15 mL falcon tube. Cells were pelleted, re-suspended in 10 mL of complete media and allowed to recover for 1.5 hours at 37°C with gentle shaking. Following cell recovery step, cells were counted using Trypan blue and re-suspended to densities between 2^6 and 2.67^6 cells/mL in assay buffer. 45 μl/well of this cell solution was added to the plate containing 15 μl/per well of inhibitor using an Agilent Bravo liquid handler. Final cell concentrations were between 1.5^6 and 2^6 cells/mL and final inhibitor concentrations were in a range from 0 to 200 nM. Cells were incubated with the inhibitors for 1 hour at RT with gentle rocking, after which 50 μL per well was transferred to the Epic assay plate, and a time course of cell adhesion was monitored for an additional 3 hours.

### Latent TGFβ activation assay

Transfected mink lung epithelial cell (TMLC) cell line containing the TGFβ responsive PAI-1 promoter driving luciferase expression ^28^ and the CHO-K1 human and mouse αvβx integrin cell lines were grown under optimized growth conditions to 90% confluency. Cells were trypsinized and plated at 25,000 cells/well in 50 μl in Costar clear bottom white plates. Dose response curve of the antibodies was added in 25 μl (in 4X concentrations) to wells containing 25,000 cells/25 μl of human or mouse αvβx cells. The mixture was transferred to the TMLC cells plated in clear bottom white Costar plates for the total volume of 100 μl and incubated at 37°C for 16 hours. 100 μl of ONE-Glo kit (#PRE6120) was added to each well and read after 3-5 minutes on the Envision.

### αSMA imaging

Primary cells (passage 3-5) were grown on Cell carrier Ultra, col1 coated 96-well plates (Perkin Elmer, #6055700) at a density of 25,000 cells/well. Once the cells adhere, they were starved in serum free media for 24 hours followed by addition of treatments i.e. 5 ng/ml TGFβ (Biolegend #580702) ± inhibitors. Cells were pre-treated with inhibitors for 30 minutes prior to the addition of TGFβ stimulation. 48hours post treatment, the cells were fixed in 4% methanol free paraformaldehyde (ThermoFisher Scientific #28908) for 20 minutes at room temperature and incubated with anti-alpha smooth muscle Actin antibody (Abcam #ab7817) at 1:4000 overnight at 4°C. Following primary Ab incubation, cells were incubated with Goat anti-mouse secondary antibody (ThermoFisher Scientific #A32723) at 1:500 and Hoechst (ThermoFisher Scientific #62249) at 1:1000 for 1 hour at room temperature. The fixed and stained cells were images for immunofluorescence on Perkin Elmer’s Opera Phenix high content Imager.

### Mouse Bleomycin Lung Fibrosis Models

Adult male C57BL/6 mice (Taconic, Rensselaer, NY) were housed in a temperature and humidity-controlled facility with a 12 hour light: 12-hour dark cycle. Animals had ad libitum access to food (Purina Rodent Chow 5053, LabDiet, St. Louis, MO) and water. All procedures utilizing experimental animals were conducted in accordance with the Guide for the Care and Use of Laboratory Animals, and experimental procedures were reviewed and approved by the Institutional Animal Care and Use Committee at MRL, Kenilworth, NJ.

For osmotic minipump implantation, TFA salt of MK-0429 was formulated in 50% DMSO/50% H_2_O at a concentration of 416 mg/ml. MK-0429 or vehicle solution were filled in minipumps (Alzet, # 1007D, flow rate 0.5 μl/hour). Minipumps were placed in mice subcutaneously in a pocket on the back between the shoulder blades. A small incision was made and a subcutaneous pocket formed by blunt dissection. A sterile minipump was inserted into the subcutaneous pocket. The incision was closed with staples or non-absorbable suture or absorbable (subcuticular) suture. The minipumps were used for continuous drug delivery for 1 week.

Mice at 12~13 weeks of age were randomized to 5 groups: saline, bleomycin instillation with vehicle (osmotic minipump), bleomycin instillation with MK-0429 treatment, bleomycin instillation with vehicle oral treatment, and bleomycin instillation with Nintedanib treatment. All mice were anaesthetized with isoflurane. Bleomycin was dosed by intra-tracheal (i.t) instillation in a volume of 50 μl, at a dose of 1unit /kg body weight. After instillation, mice were kept in a heads-up position for 2-5 minutes before putting into cages. MK-0429 (200 mg/kg) was administered by Osmotic pump from day 5 to day 14 and Nintedanib (60 mg/kg) was administered, p.o., q.d., for 14 days. Vehicle was dosed orally at 10 am daily from the day 5 to the end of the studies and dosing volume was 10 ml/kg. For plasma PK analysis, an aliquot of 50 μL plasma was spiked into a 96-well plate, and 200 μL of acetonitrile containing internal standard were added for protein precipitation. The mixture was vortexed, centrifuged at 4000 rpm for 20 minutes. 50 μL of supernatant were mixed with 200 μl H_2_O and the final solution were injected for LC-MS/MS analysis. The methods for quantitative analysis were developed on UPLC (Waters) chromatographic system equipped with an API4000 QTrap mass spectrometer (Applied Biosystems, Concord, Ontario, Canada). Analyst 1.5 software packages (Applied Biosystems) were used to control the LC-MS/MS system and data acquisition and processing.

### Histopathology Analysis

Upon completion of the study, animals were euthanized and tissues were collected for histological assessment. Lung tissues were perfused with 10% formalin, fixed for 24 hours, and then paraffin embedded. Tissue sections were stained with hematoxylin & eosin (H&E) and Masson’s trichrome, subsequently evaluated under light microscope. The severity of histopathologic changes and fibrosis in the lung were graded as described previously by pathologists ^53,54^. Following deparaffinization and rehydration, each lung tissue section was processed to identify αSMA deposition by immunohistochemistry. The primary antibodies used were αSMA antibody from Sigma. The Aperio ScanScope XT Slide Scanner (Aperio Technologies) system was used to capture whole slide digital images with a 20× objective. Digital images were managed using Aperio Spectrum. The positive stains were identified and quantified using a macro created from a color deconvolution algorithm (Aperio Technologies). Statistical analysis was performed by using One-way ANOVA followed by Tukey’s test.

### Integrin co-immunoprecipitation and western blot analyses

Cells or lung tissue were lysed with assay buffer (50mM Tris, pH7.4, 150mM NaCl, 1mM EDTA, 10% glycerol, 2% NP-40) with the addition of protease inhibitor (Roche cOmplete mini-pellet, EDTA free, #4693159001) and phosphatase inhibitor cocktails (Sigma #P5726). The total protein concentration was determined by Bradford reagent (Bio-Rad #5000006). 1mg of cell lysates were incubated with anti-αv antibody (Enzo #ALX-803-304-C100) and Protein G magnetic beads (Pierce #88802) at 4°C overnight. After washes, the beads were treated with 0.1X sample buffer with fluorescent dye and boiled at 95°C for 5 minutes. The supernatant was subsequently subjected for Sally Sue Simple Western analysis from ProteinSimple ^55^. The antibodies used for detecting integrin isoforms were shown in **Supplemental Table S2**, and anti-FAK and GAPDH antibodies were from Cell Signaling Transduction.

### Molecular modeling

To generate a model for our antibodies, we employed the Antibody Modeler tool within MOE 2019 (Chemical Computing Group, Montreal, Canada) using the determined sequence for Ab-31 and related antibodies to generate a final Fab model. Default settings in MOE 2019 were used during model generation, with the highest scoring model of the ten produced models selected for further study.

To model how our antibodies may bind to integrin, previously published structures of α5β1 integrin (PDB: 3VI4) and αvβ3 (PDB: 6AVQ) were used as models for antibody binding to integrin ^37,56^. Each structure was first prepared by deleting additional copies of integrin in the original crystal structures, followed by running the Structure Preparation and Protonate 3D tools within MOE 2019 to add missing side chains, loops, hydrogens, and cap termini using the Amber14EHT force field. Protein-Protein docking was then carried out in MOE 2019 using integrin as the receptor and the modeled antibody Fv region as the ligand while enabling both hydrophobic patch potentials and restraining the ligand site to the CDRs of the modeled Ab-31 structure as annotated by MOE 2019. The resulting poses were then visualized across Ab-31 and related antibodies which were also modeled, as well as across two distinct integrin structures to identify poses that were consistent with the previously described experimental data. The final model was then compared the published integrin-antibody complexes of Abituzumab (PDB: 4O02) and LM609 (PDB: 6AVQ) ^39,43^. All images were generated in MOE 2019.

## Supplemental Information

### Supplemental Figures

**Supplemental Fig. S1.**
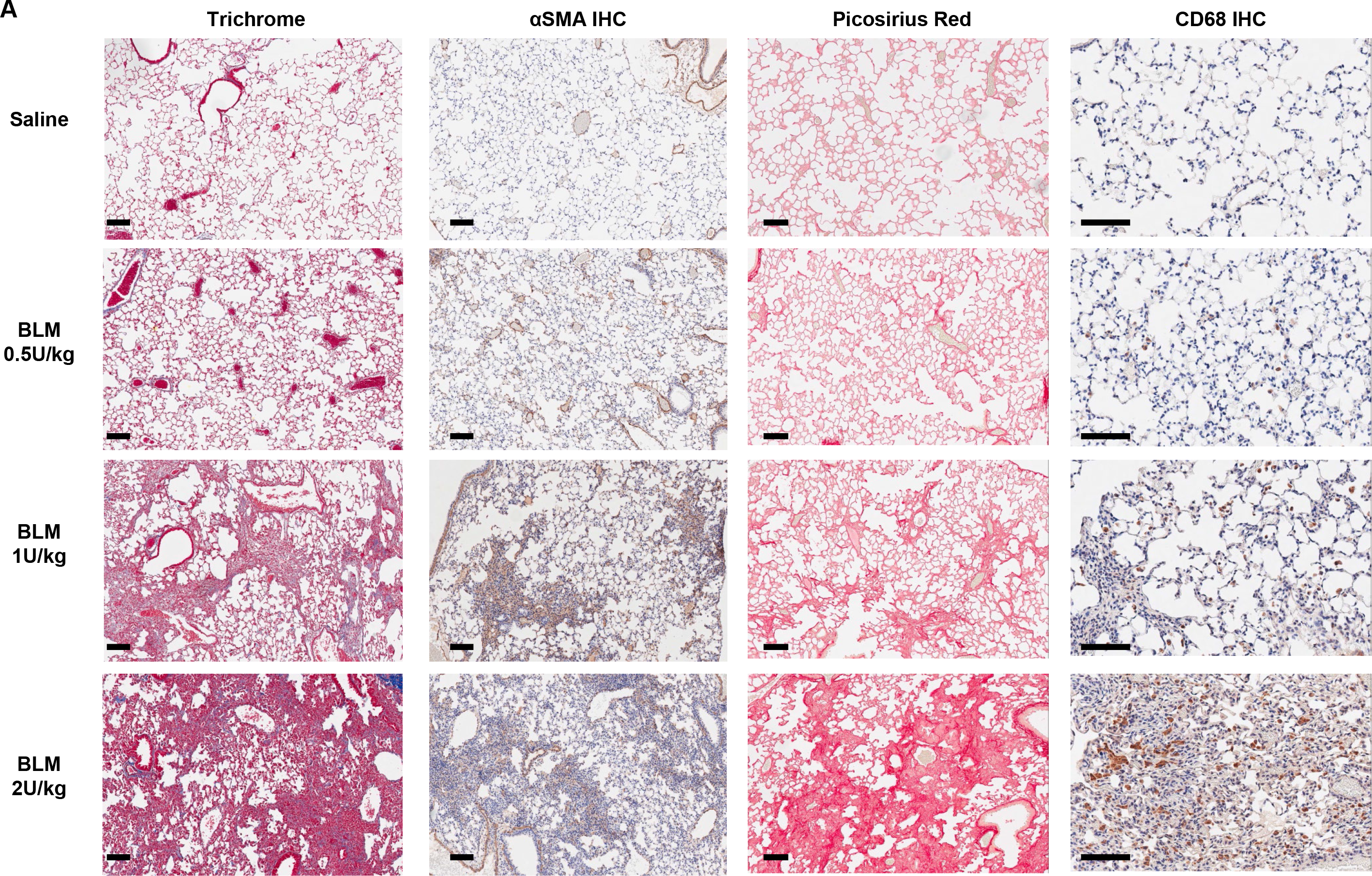
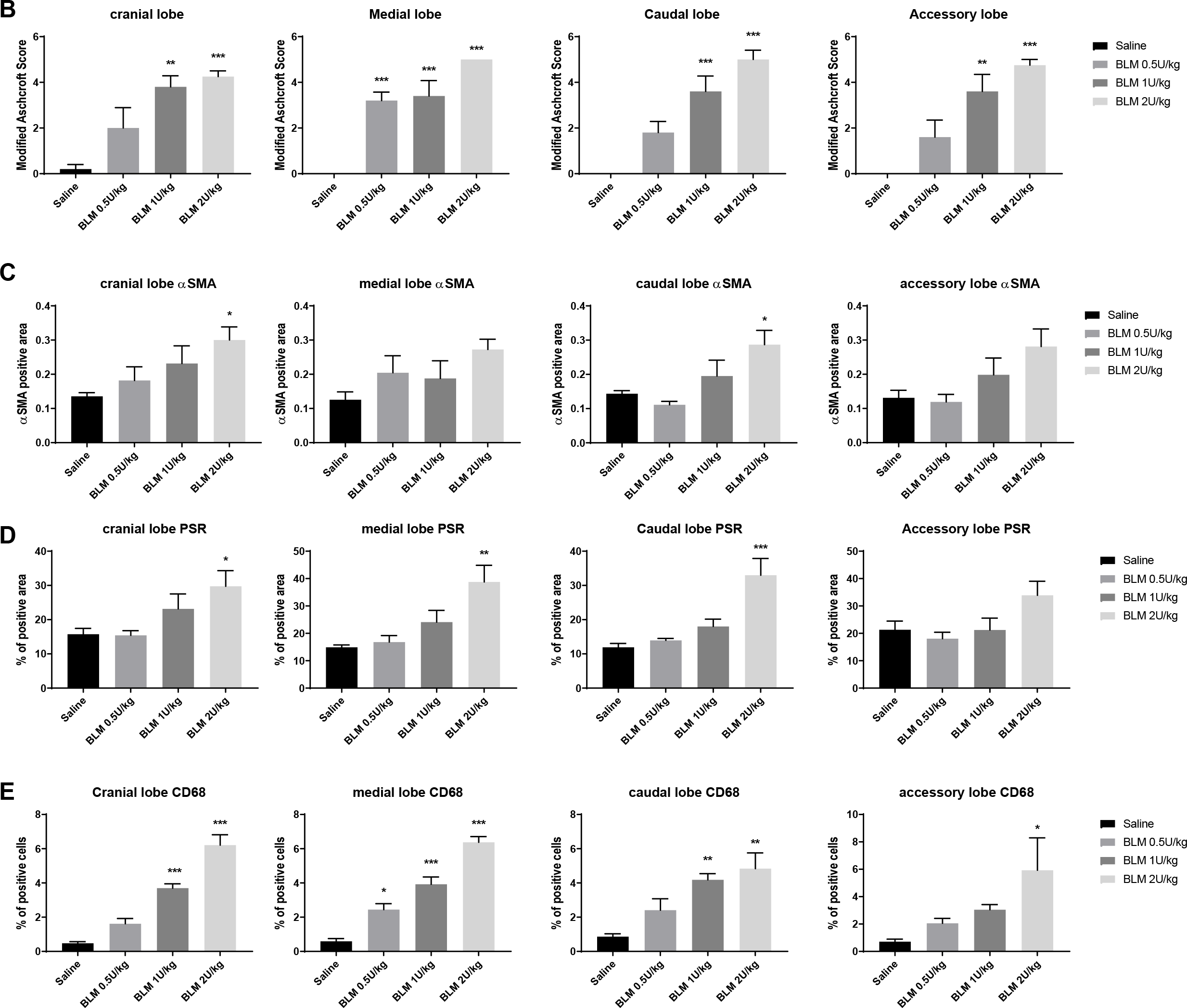
Histology analysis of bleomycin-induced lung fibrosis model. A) Representative histological images of mouse lungs from each experimental group. Scale bar, 100 μM. Immunohistochemistry, IHC. Quantitative analysis of each lung lobe, B) Modified Ashcroft score, C) αSMA positive area, D) Area of Picosirus red (PSR) positive staining, and E) Percentage of CD68-positive cells. Mean±SEM, n=5. One-way ANOVA followed by Tukey’s test, *p<0.05, **p<0.01, ***p<0.005 vs Saline group.

**Supplemental Fig. S2.**
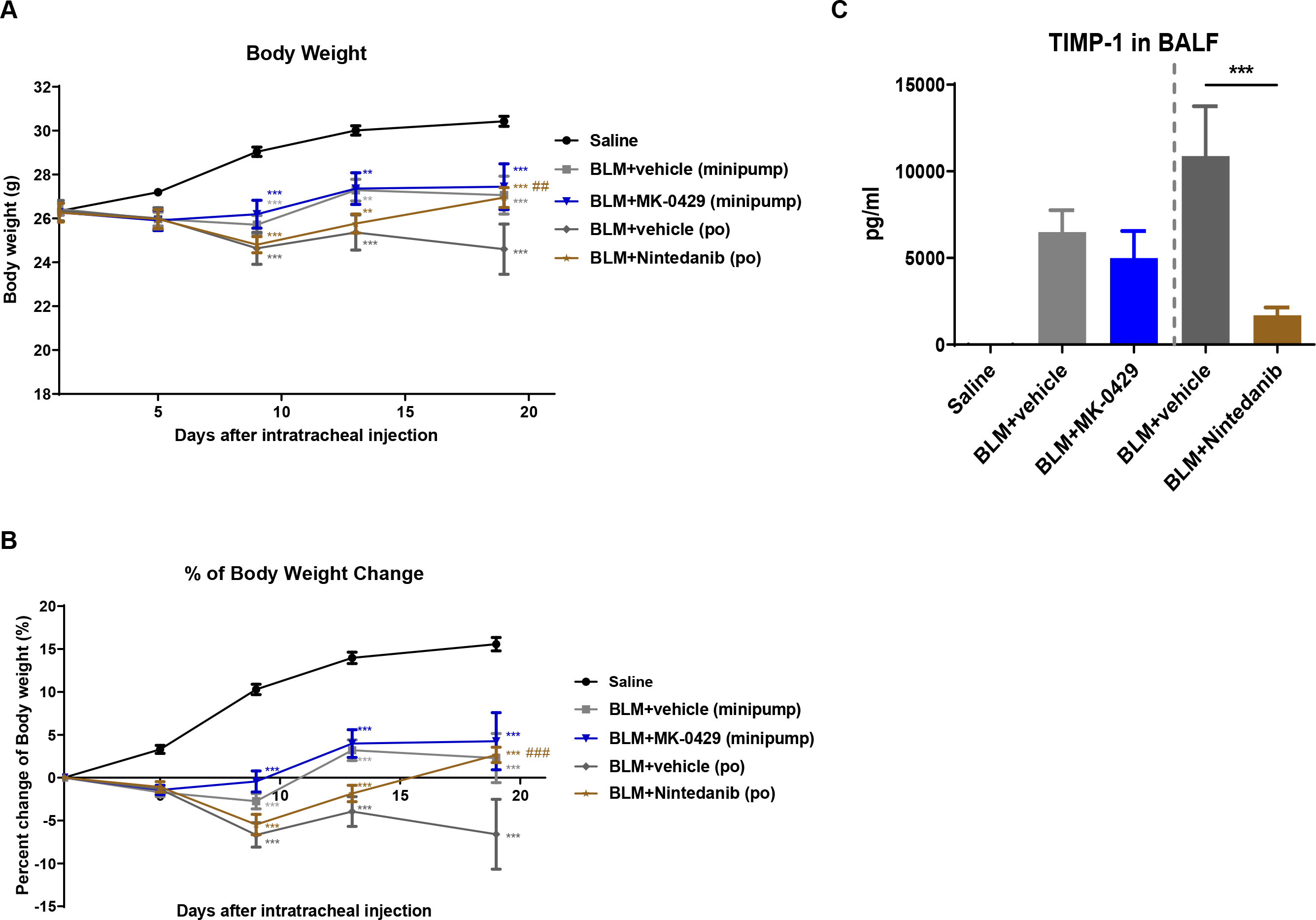
MK-0429 inhibits lung fibrosis in mouse bleomycin model. A) Body weight of each experimental group over the duration of time course. Mean±SEM, n=10. One-way ANOVA followed by Tukey’s test, **p<0.01, ***p<0.001 vs Saline group; ##p<0.01, ###p<0.001 vs BLM-vehicle group. B) Percentage of body weight changes in each experimental group. Mean±SEM, n=10. One-way ANOVA followed by Tukey’s test, **p<0.01, ***p<0.001 vs Saline group; ##p<0.01, ###p<0.001 vs BLM-vehicle group. C) TIMP1 concentration in BALFs. Mean±SEM, n=10. One-way ANOVA followed by Tukey’s test, *p<0.05, **p<0.01, ***p<0.001 vs BLM-vehicle group.

**Supplemental Fig. S3.**
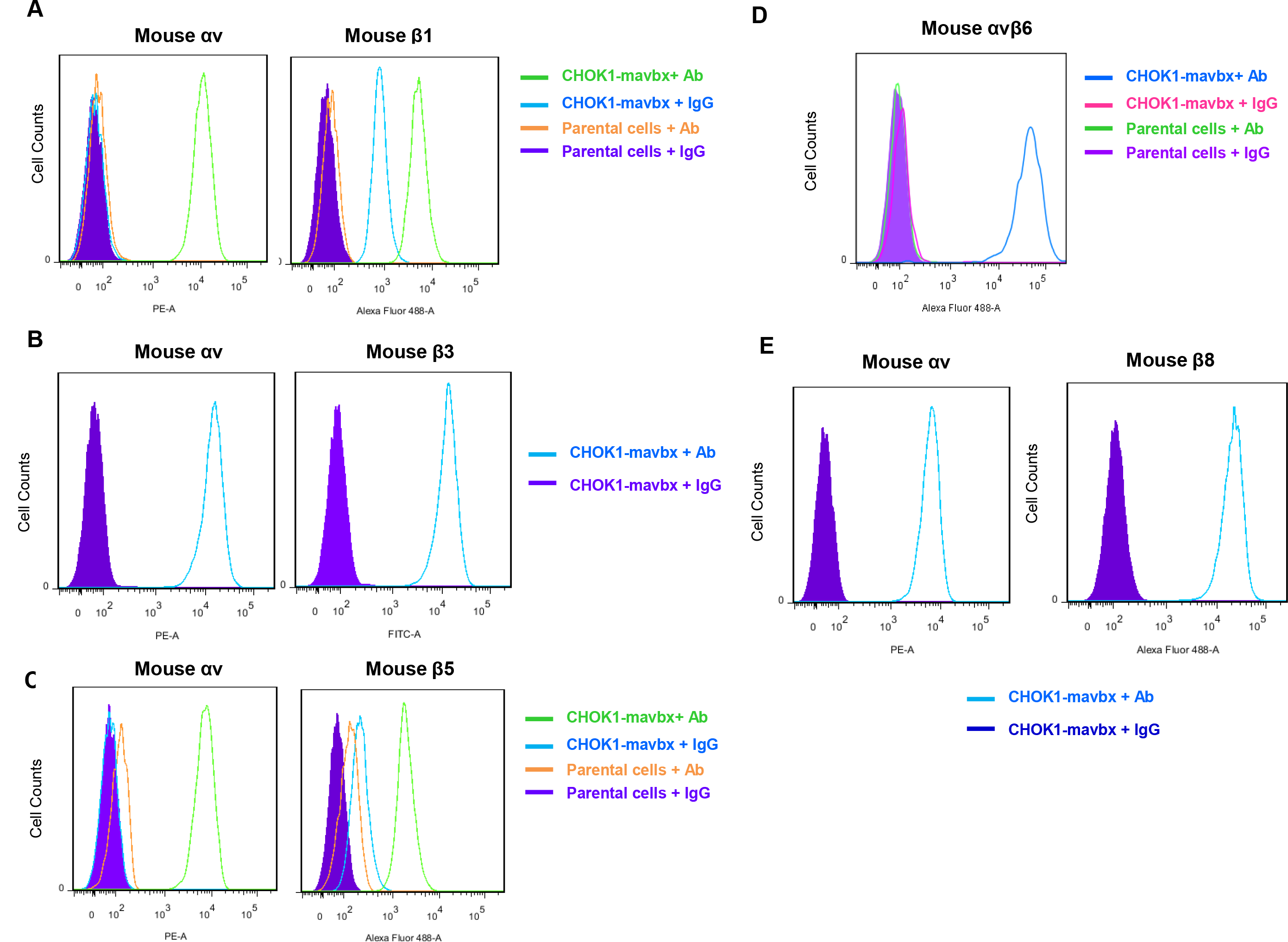
The expression of αv integrins in CHOK1 stable lines. A) FACS of mouse αv and β1 expression in CHOK1-α5KO-mαvβ1 cells. Anti-αv (RMV7) antibody and anti-β1 (KMI6) antibody were used for detection. B) FACS of mouse αv and β3 expression in CHOK1-mαvβ3 cells. Anti-αv (RMV7) antibody and anti-β3 (HMβ3.1) antibody were used for detection. C) FACS of mouse αv and β5 expression in CHOK1-mαvβ5 cells. Anti-αv (RMV7) antibody and anti-β5 (P1F76) antibody were used for detection. D) FACS of mouse αvβ6 expression in CHOK1-mαvβ6 cells. Anti-αvβ6 (10D5) antibody was used for detection. E) FACS of mouse αv and β8 expression in CHOK1-mαvβ8 cells. Anti-αv (RMV7) antibody and anti-β8 (ADWA11) antibody were used for detection.

**Supplemental Fig. S4.**
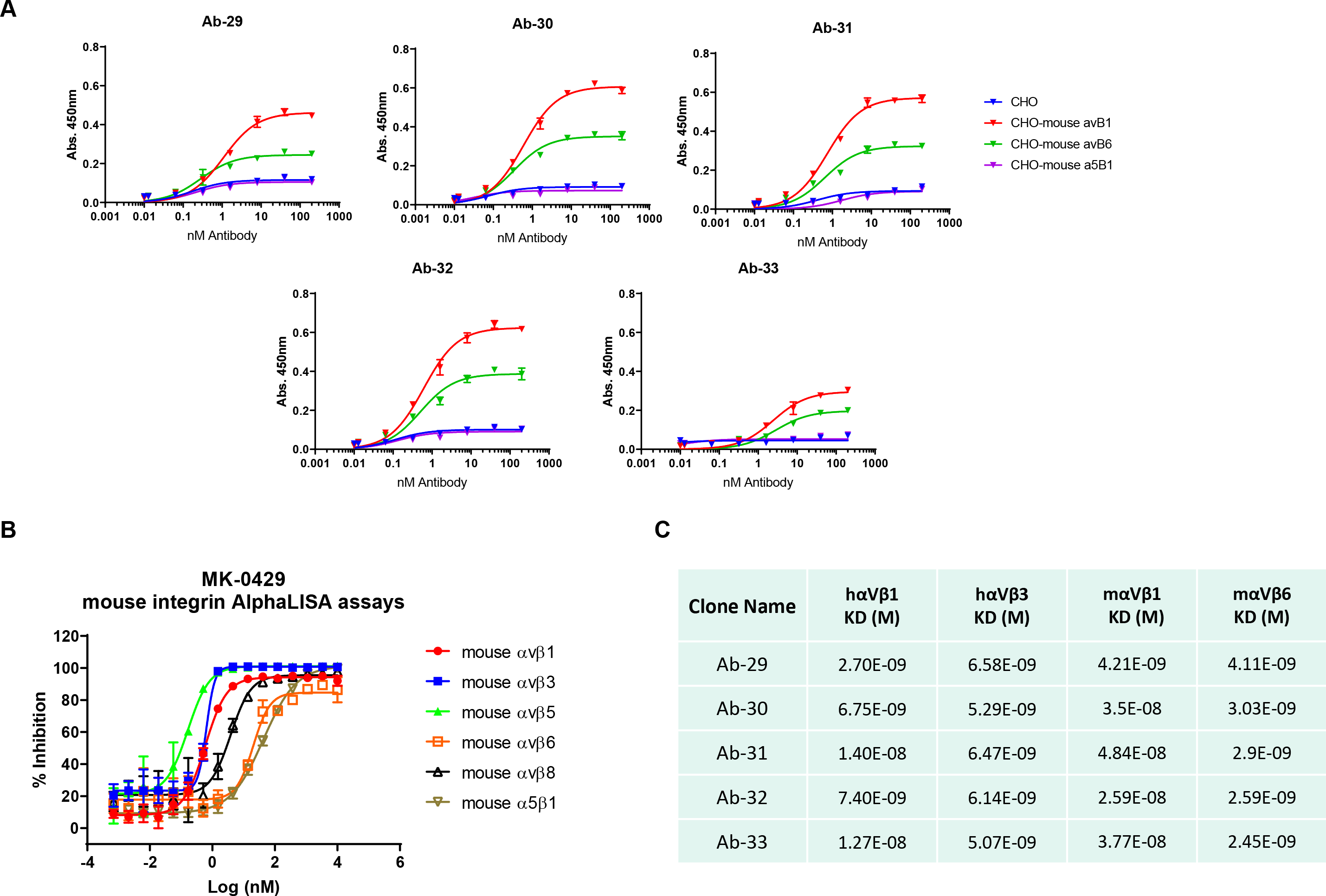
Integrin antibodies with strong blocking activities against both human and mouse αv integrins. A) A) Titration of Ab-29, Ab-30, Ab-31, Ab-32, and Ab-33 for their binding to CHOK1-mouse αvβ1, αvβ6, and α5β1 stable cell lines in CELISA assays. B) Dose-dependent inhibition of mouse integrin-ligand binding by MK-0429 in AlphaLISA assay panel. C) Octet Red binding parameter of Ab-29, Ab-30, Ab-31, Ab-32, and Ab-33 to selected human and mouse integrins.

**Supplemental Fig. S5.**
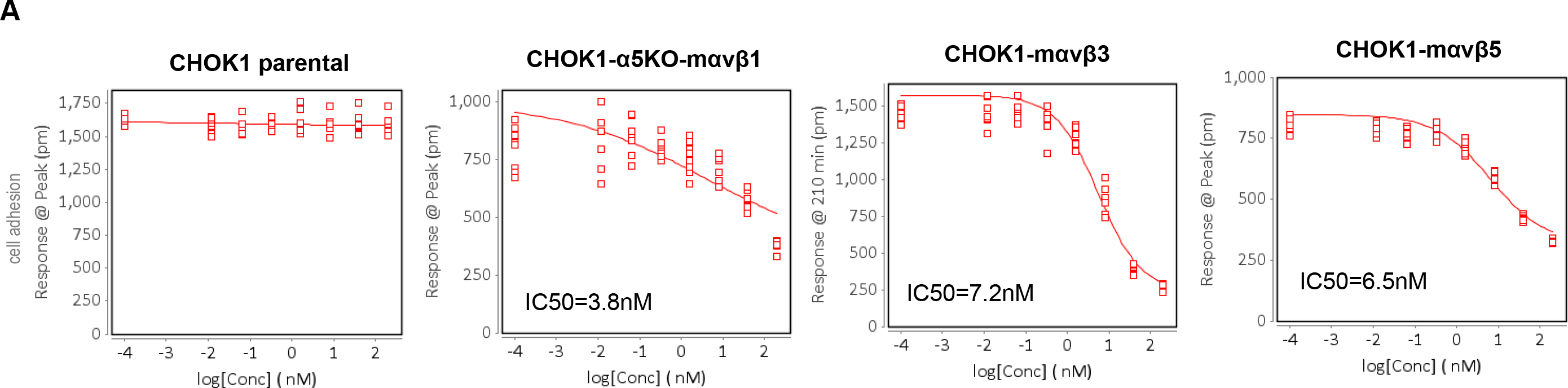
MK-0429 inhibits integrin-mediated cell adhesion. A) The effect of MK-0429 on the adhesion of CHOK1 parental, CHOK1-α5KO-mαvβ1, CHOKl-mαvβ3, and CHOK1-mαvβ5 cells to fibronectin or vitronectin matrix.

**Supplemental Fig. S6.**
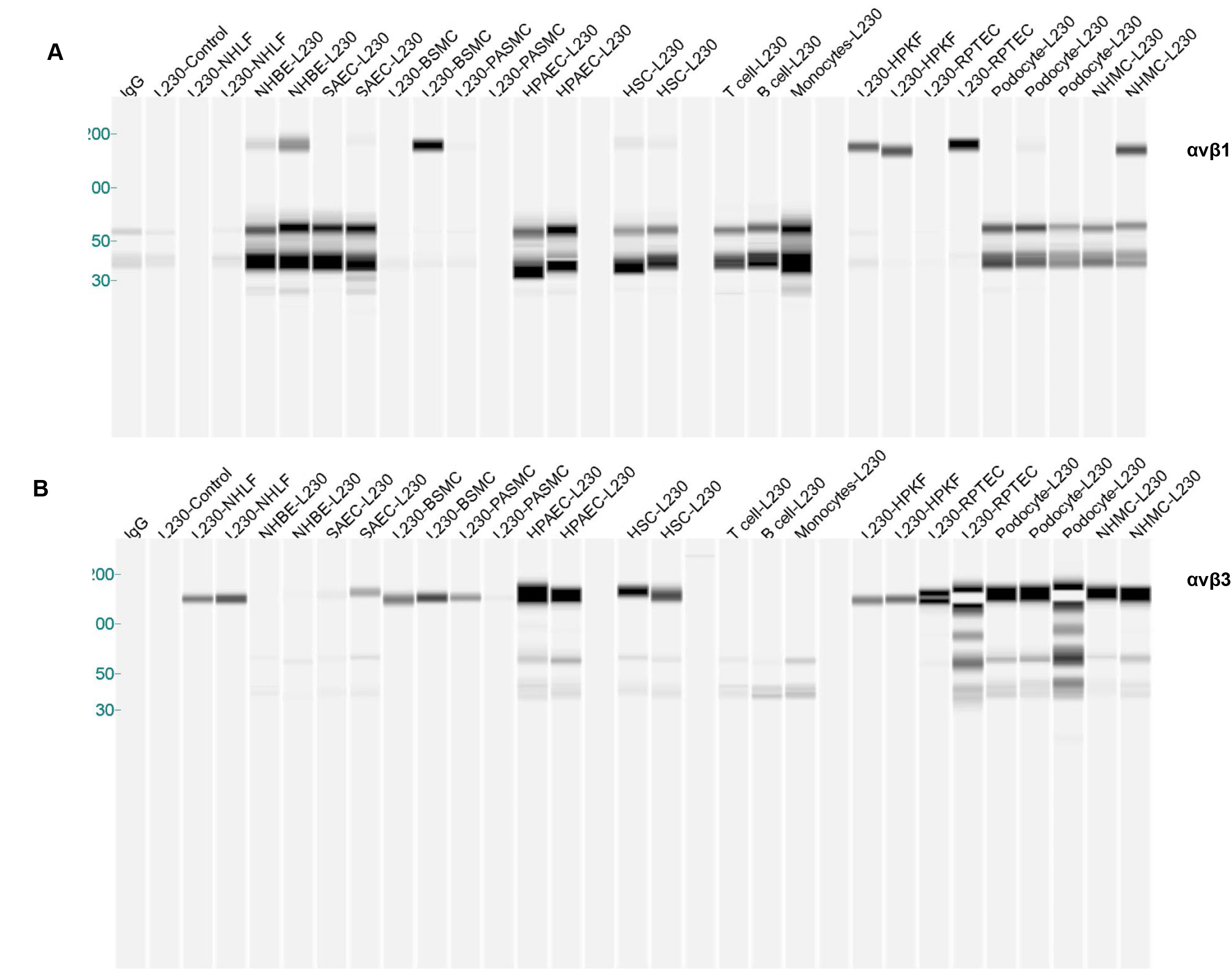
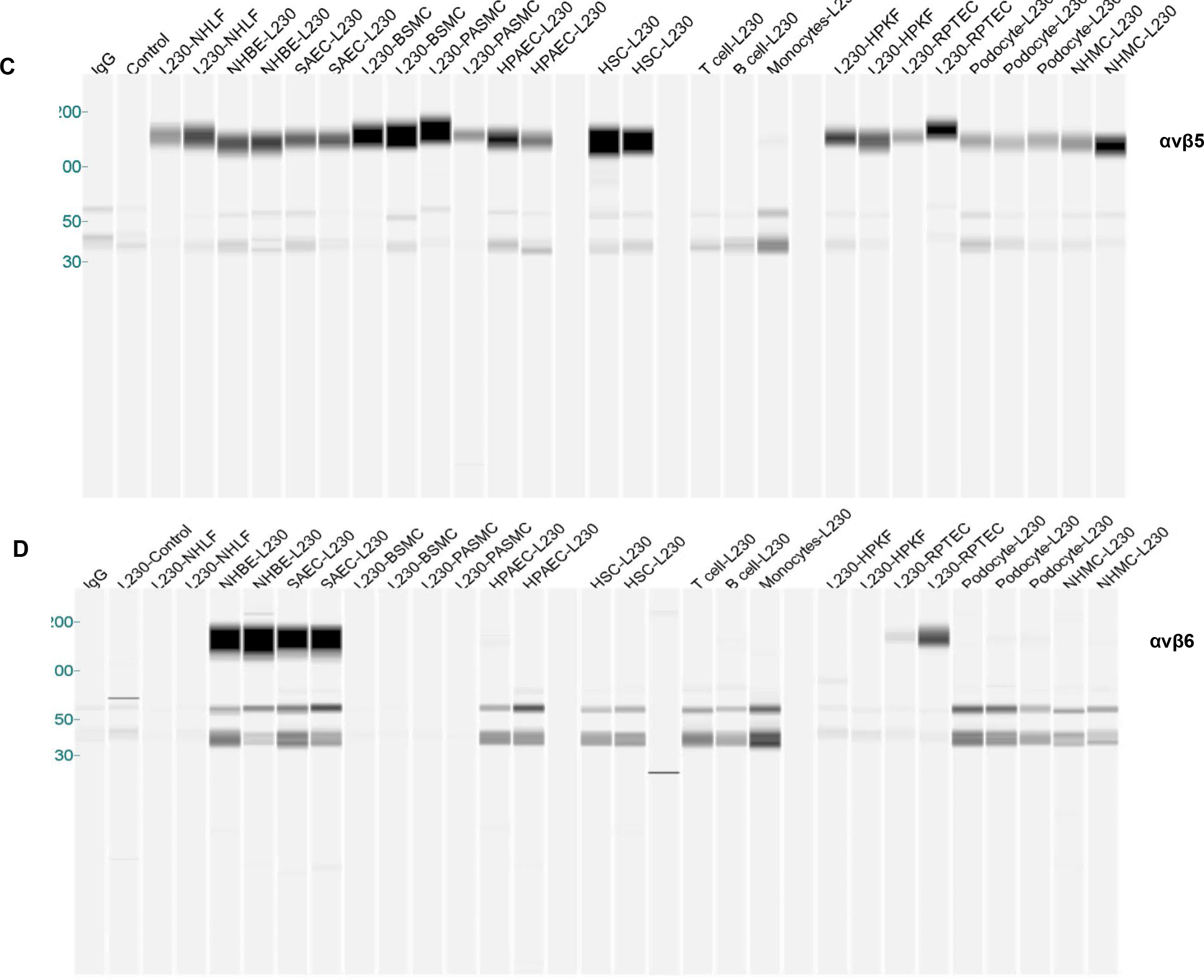
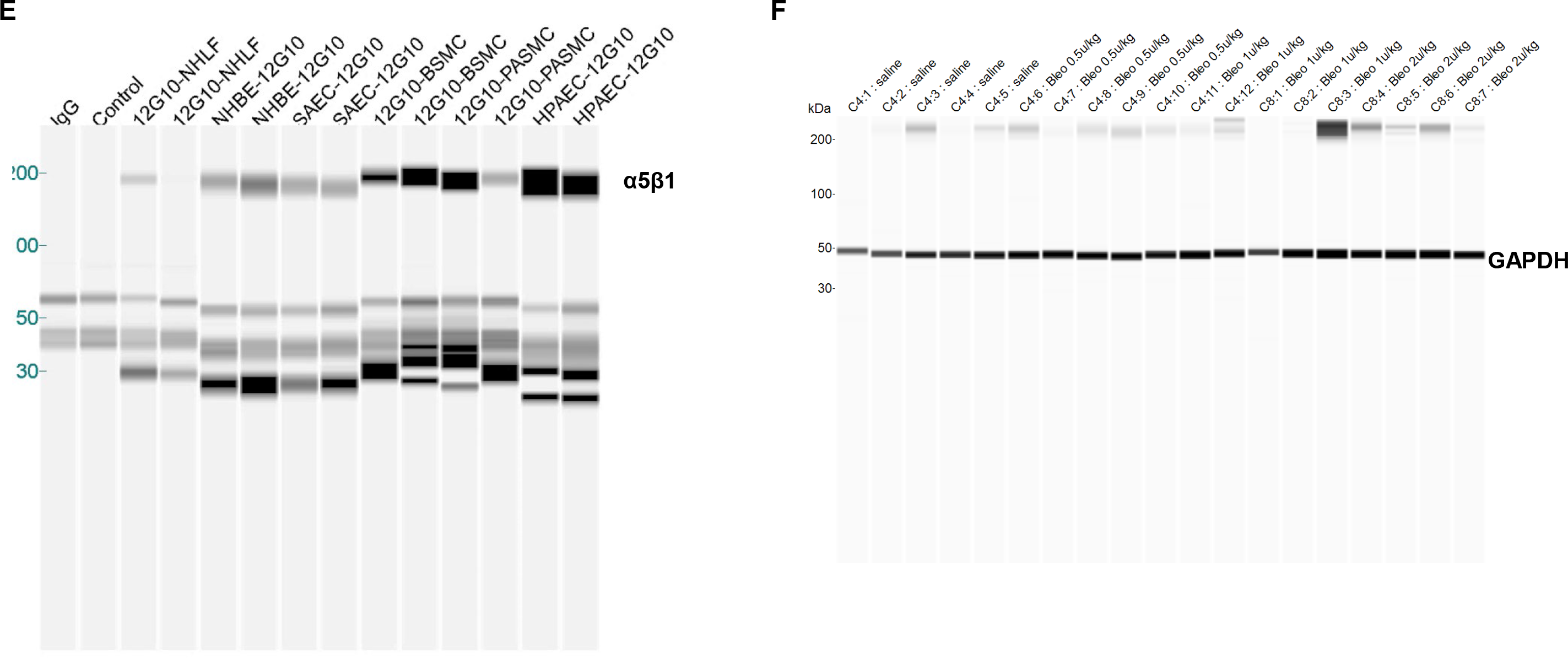
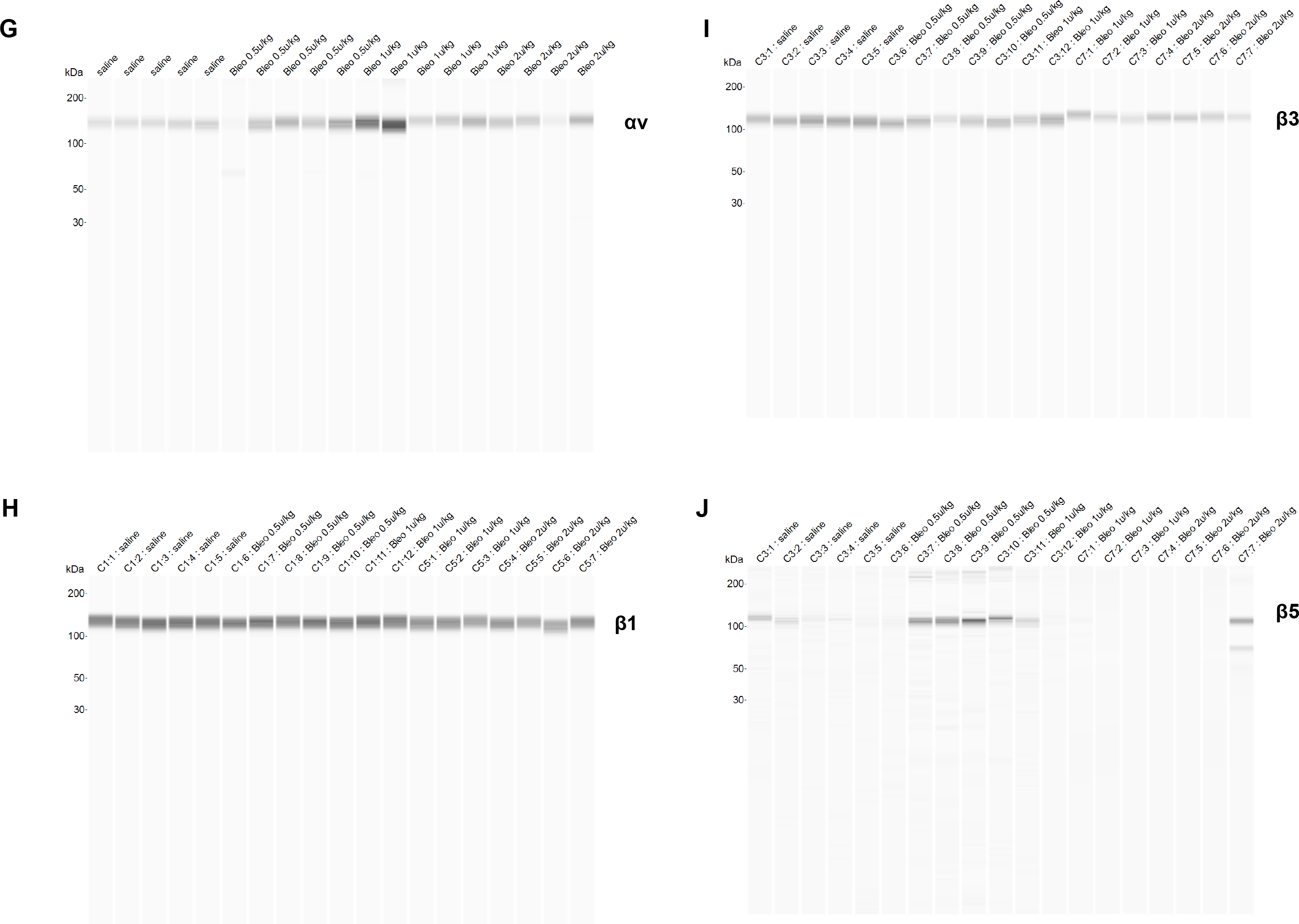
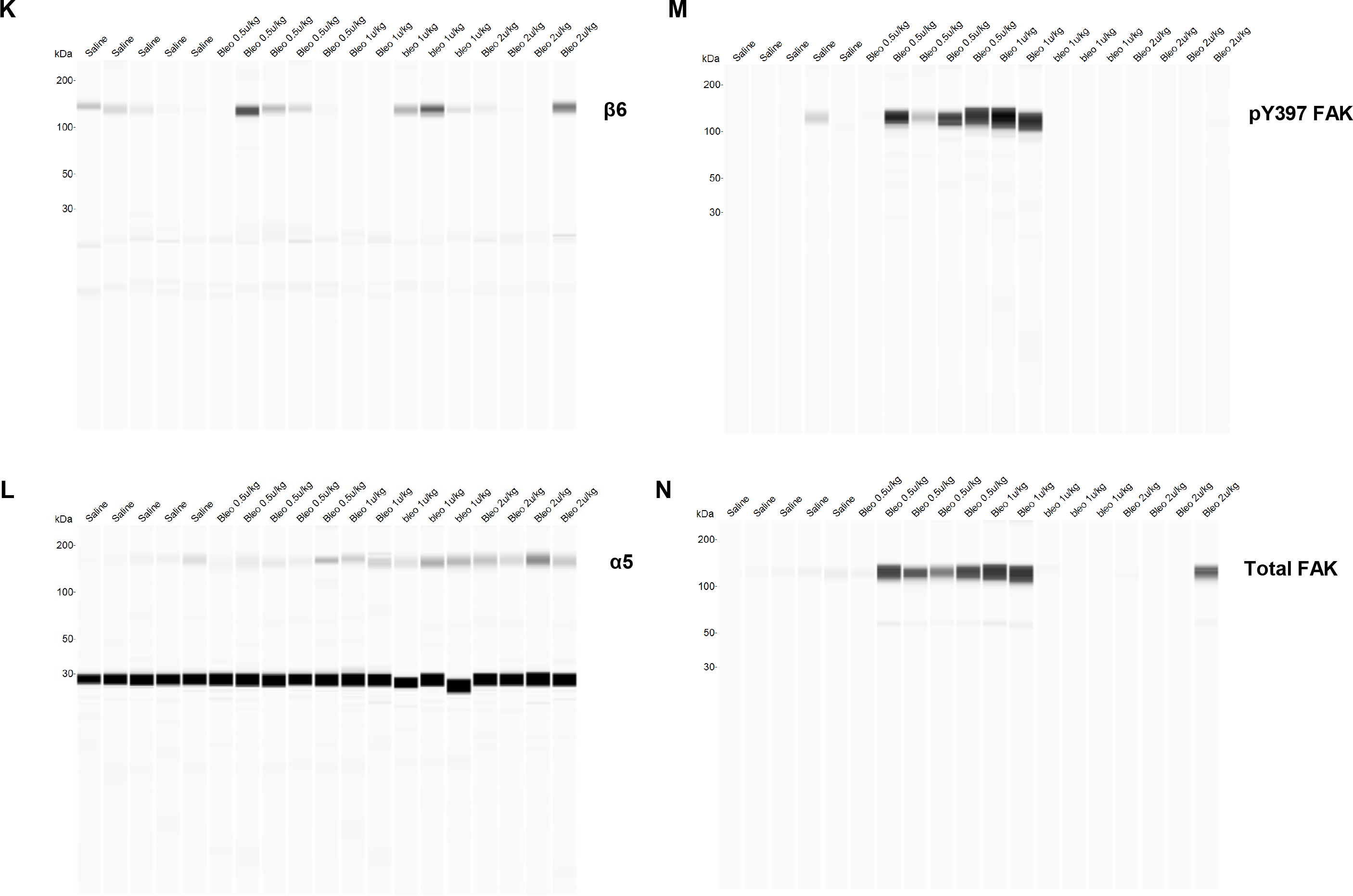
Sally Sue simple western full-length blot images presented in Fig. 1. A-E) blot images presented in Fig. 1A. The expression of various integrins in human primary lung cell types upon TGFβ (5ng/ml for 24 hours) treatment. Following immunoprecipitation with an anti-αv antibody, the αvβ1, αvβ3, αvβ5, and αvβ6 heterodimers were detected by Sally Sue simple western analysis after using antibodies that recognize each individual β-subunit. Normal human lung fibroblast, NHLF; normal human bronchial epithelial cells, NHBE; small airway epithelial cells, SAEC; bronchial smooth muscle cells, BSMC; pulmonary artery smooth muscle cells, PASMC; pulmonary artery endothelial cells, PAEC. Integrin antibodies were reported in Supplemental Table S2. Exposure time: 4 seconds. F-N) blot images presented in Fig. 1C. Integrin expression and signaling in fibrotic lungs was determined by Sally Sue simple western analysis using antibodies that recognized the individual subunits. GAPDH level in total lung lysates was used as a loading control. Integrin antibodies were reported in Supplemental Table S2. Exposure time: 4 seconds.

**Supplemental Table S1.** Recombinant human and mouse integrin expressing constructs.

| **Integrin subunit** | **N-term** | **C-term** |
| --- | --- | --- |
| hαV | 1 | 992 |
| hα5 | 1 | 995 |
| hβ1 | 1 | 728 |
| hβ3 | 1 | 718 |
| hβ5 | 1 | 719 |
| hβ6 | 1 | 709 |
| hβ8 | 1 | 684 |
| mαV | 1 | 988 |
| mα5 | 1 | 999 |
| mβ1 | 1 | 728 |
| mβ3 | 1 | 717 |
| mβ5 | 1 | 719 |
| mβ6 | 1 | 706 |
| mβ8 | 1 | 679 |

**Supplemental Table S2.** Integrin antibodies used for FACS and Sally Sue simple western.

| **Integrins** | **Clone** | **Company** | **Catalog #** | **Isotype** | **Reactivity** |
| --- | --- | --- | --- | --- | --- |
| $\alpha 5$ | HMα5-1 | BD | 553350 | hamster IgG1 | mouse |
| $\alpha 5$ | EPR7854 | Abcam | 150361 | rabbit IgG | human, mouse |
| $\alpha 5$ | PB1 | DSHB | | mouse IgG1 | hamster |
| $\alpha v$ | RMV-7 | BD | 552299 | rat IgG1 | mouse |
| $\alpha v$ | P2W7 | R&D Systems | MAB1219 | mouse IgG1 | human |
| $\beta 1$ | KMI6 | BD | 558741 | rat IgG2a | mouse |
| $\beta 1$ | P5D2 | R&D Systems | MAB17781 | mouse IgG1 | human |
| $\beta 1$ | 7E2 (DSHB) | DSHB | | mouse IgG1 | hamster |
| $\beta 3$ | HMβ3.1 | BD | 553347 | hamster IgG1 | |
| $\alpha v\beta 3$ | 27.1(VNR-1) | Abcam | ab78289 | mouse IgG1 | human |
| $\beta 5$ | P1F6 | Millipore | MAB1961 | mouse IgG1 | mouse |
| $\alpha v\beta 5$ | P5H9 | R&D Systems | MAB2528 | mouse IgG1 | human |
| $\alpha v\beta 6$ | 10D5 | BD | 566922 | mouse IgG2a | human, mouse |
| $\beta 8$ | ADWA11 | - | - | mouse IgG1 | human, mouse |
| $\beta 8$ | 416922 | LSBio | LS-C70803 | mouse IgG2b | human |

**Supplemental Table S3.** AlphaLISA assay condition and reagents.

| Human AlphaLISA Integrin Assays: Final Conditions |  |  |  |  |  |  |  |  |  |  |  |  |  |  |
| --- | --- | --- | --- | --- | --- | --- | --- | --- | --- | --- | --- | --- | --- | --- |
| Integrin EC80<br>(batch dependent) |  |  | Ligand |  |  | Acceptor Bead Solution |  |  |  |  |  | Donor |  |  |
|  |  |  |  |  |  | antibody |  |  | AlphaLISA Acceptor |  |  |  |  |  |
| havb1 | 1.4 | nM | human Fibronectin<br>R&D Systems (1918-FN) | 1.0 | nM | n/a | | | $\alpha$ - Fibronectin<br>Acceptor beads/ Perkin<br>Elmer (CUSM03822000)<br>See notes | 10.0 | ug/ml | Streptavidin Donor<br>beads/ Perkin Elmer<br>(6760002) | 10.0 | ug/ml |
| ha5b1 | 0.4 | nM |  | 0.5 | nM |  |  |  |  |  |  |  |  |  |
| havb3 | 3.0 | nM | human Vitronectin-GST<br>tagged<br>EMD Millipore (08-126) | 10.0 | nM | n/a | | | $\alpha$ - GST Acceptor<br>beads/ Perkin<br>Elmer(AL110C) | 15.0 | ug/ml | | 15.0 | ug/ml |
| havb5 | 2.5 | nM |  | 1.0 | nM |  |  |  |  |  |  |  |  |  |
| havb6 | 0.1 | nM | human LAP<br>R&D system (246-LP) | 0.5 | nM | α-Human LAP (goat<br>pAb) R&D system (AF-246-NA) | 3.0 | nM | $\alpha$ - goat IgG Acceptor<br>beads/ Perkin Elmer<br>(AL107C) | 15.0 | ug/ml | | 15.0 | ug/ml |
| havb8 | 1.0 | nM |  | 3.0 | nM |  | 3.0 | nM |  |  |  |  |  |  |
| Mouse AlphaLISA Integrin Assays: Final Conditions |  |  |  |  |  |  |  |  |  |  |  |  |  |  |
| Integrin EC80<br>(batch dependent) |  |  | Ligand |  |  | Acceptor Bead Solution |  |  |  |  |  | Donor |  |  |
|  |  |  |  |  |  | antibody |  |  | AlphaLISA Acceptor |  |  |  |  |  |
| maVb1 | 4.3 | nM | mouse fibronectin Abcam<br>(ab92784) | 1.0 | nM | $\alpha$ - Fibronectin (Rabbit<br>pAb) Abcam(ab2413) | 5.0 | nM | α-rabbit IgG Acceptor/<br>Perkin Elmer (AL104C) | 15 | ug/ml | Streptavidin Donor<br>beads/ Perkin Elmer<br>(6760002) | 15 | ug/ml |
| ma5b1 | 1.0 | nM |  | 2.5 | nM |  |  |  |  |  |  |  |  |  |
| maVb3 | 3.0 | nM | mouse Vitronectin Abcam<br>(ab92727) | 5.0 | nM | $\alpha$ - Vitronectin (Rabbit<br>pAb) Abcam<br>(ab140016) | 2.0 | nM | | | | | | |
| maVb5 | 0.6 | nM |  | 2.5 | nM |  |  |  |  |  |  |  |  |  |
| maVb6 | 1.4 | nM | mouse LAP-6His<br>Chempartner | 2.5 | nM | n/a | | | α-6xHIS/ Perkin Elmer<br>(AL128) | 15 | ug/ml | | 15 | ug/ml |
| maVb8 | 1.0 | nM |  | 2.5 | nM |  |  |  |  |  |  |  |  |  |

